# HSV-1 infection induces a downstream shift of promoter-proximal pausing for most host genes

**DOI:** 10.1101/2022.09.28.509911

**Authors:** Elena Weiß, Thomas Hennig, Pilar Graßl, Lara Djakovic, Adam W. Whisnant, Christopher S. Jürges, Franziska Koller, Michael Kluge, Florian Erhard, Lars Dölken, Caroline C. Friedel

## Abstract

Herpes simplex virus 1 (HSV-1) infection exerts a profound shut-off of host gene expression at multiple levels. Recently, HSV-1 infection was reported to also impact promoter-proximal RNA Polymerase II (Pol II) pausing, a key step in the eukaryotic transcription cycle, with decreased and increased Pol II pausing observed for activated and repressed genes, respectively. Here, we demonstrate that HSV-1 infection induces more complex alterations in promoter-proximal pausing than previously suspected for the vast majority of cellular genes. While pausing is generally retained, it is shifted to more downstream and less well-positioned sites for most host genes. We exclude that this is an artefact of alternative *de novo* transcription initiation at downstream sites or read-in transcription originating from disruption of transcription termination for upstream genes. Use of downstream 2^nd^ pause sites associated with +1 nucleosomes was previously observed upon NELF depletion. However, downstream shifts of Pol II pausing upon HSV-1 infection are much more pronounced than observed upon NELF depletion. Thus, our study reveals a novel aspect in which HSV-1 infection fundamentally reshapes host transcriptional processes with implications for our understanding of maintenance of promoter-proximal Pol II pausing in eukaryotic cells.

## Introduction

Lytic Herpes simplex virus 1 (HSV-1) infection exerts a profound shutoff of host gene expression via degradation of host and viral mRNAs mediated by the HSV-1 virus host shutoff protein (*vhs)* (1, 2) and inhibition of host transcriptional activity (3–6). Efficient recruitment of RNA polymerase II (Pol II) and elongation factors from the host chromatin to replicating viral genomes leads to a substantial loss of Pol II occupancy from the host genome as early as 2-3 h post infection (h p.i.) (3–6). By 8 h p.i., host transcriptional activity is estimated to be only 10-20% of uninfected cells (7). We previously showed that HSV-1 infection disrupts transcription termination for the majority but not all cellular genes, leading to read-through transcription for tens-of-thousands of nucleotides beyond the poly(A) site (8). More recently, Rivas *et al*. revealed that HSV-1 also impacts promoter-proximal pausing of Pol II on host genes (9). Following transcription initiation, Pol II pauses 20-60 nucleotides downstream of the transcription start site (TSS) (10, 11) as a consequence of one or more structural rearrangements within the transcription elongation complex (12). Pausing makes Pol II vulnerable to nucleosome-induced arrest, backtracking of the elongation complex along the DNA, and promoter-proximal premature termination (12). The elongation factor TFIIS can rescue Pol II from pausing and restart transcription by mediating cleavage of backtracked RNA (13). In contrast, 5,6-dichloro-1-beta-d-ribofuranosylbenzimidazole sensitivity-inducing factor (DSIF) and negative elongation factor (NELF) stabilize paused Pol II (14, 15). Phosphorylation of DSIF, NELF and the Pol II C-terminal domain by the CDK9 subunit of the positive transcription elongation factor b (P-TEFb) is required for the release of paused Pol II into gene bodies and the switch to productive elongation (16–18). As a consequence, inhibition of P-TEFb by CDK9 inhibitors increases promoter-proximal pausing (19). The HSV-1 immediate-early protein ICP22 inhibits Pol II transcription elongation by direct interaction with CDK9 (20) and ectopic expression of a short segment of ICP22 mimics the effects of P-TEFb inhibition on Pol II transcription (21). In wild-type HSV-1 infection, however, Rivas *et al*. found that promoter-proximal Pol II pausing was actually reduced, at least for activated genes, using Pol II ChIP-seq analysis (9). This was largely dependent on ICP4. ICP4 is one of five immediate-early proteins (including also ICP0, ICP22, ICP27, and ICP47) expressed shortly after infection and is necessary for transcription of early and late viral genes (22). HSV-1-activated genes exhibited a greater increase in Pol II occupancy on gene bodies than on promoters, consistent with increased transcriptional elongation. For repressed genes, promoter-proximal Pol II pausing was increased independently of ICP4 with Pol II occupancy decreasing more strongly on gene bodies than in the promoter region.

Here, we report on a genome-wide investigation on the impact of HSV-1 infection on promoter-proximal pausing of all expressed host genes. This is based on a re-analysis of precision nuclear run-on analysis (PRO-seq) data for mock and 3 h p.i. HSV-1 strain F (WT-F) infection from a previous study by Birkenheuer *et al*. (5). PRO-seq sequences RNA that is actively transcribed by Pol II and depicts strand-specific Pol II transcriptional activity. Transcription initiation from most human gene promoters is bidirectional with productive transcription elongation occurring only in sense direction (23–26). PRO-seq thus provides nucleotide-level resolution of Pol II activity and allows separating sense and antisense Pol II initiation and pausing. Birkenheuer *et al*. already reported that Pol II levels at the promoter-proximal pause site were altered in a gene-specific manner (5), but did not explicitly investigate Pol II pausing for host genes. In a later study, they investigated Pol II pausing for HSV-1 genes (27). Our re-analysis uncovered that HSV-1 infection appears to lead to a reduction of Pol II pausing for the majority of genes when using standard Pol II pausing analysis methods. More detailed analyses, however, revealed that Pol II pausing is retained in HSV-1 infection for most host genes but shifted to sites further downstream of the promoter. In contrast to well-defined Pol II pausing peaks at the TSS observed in mock infection, HSV-1 infection resulted in more varied and less well-positioned patterns of Pol II pausing. This included both broadening of Pol II pausing peaks into downstream regions for some genes as well as newly originating or increasing Pol II peaks at downstream sites for other genes. Delayed promoter-proximal pausing at 2^nd^ downstream pause sites associated with +1 nucleosomes has recently been reported upon NELF depletion (28). However, our analysis showed that pausing is shifted even further downstream upon HSV-1 infection. In summary, our study demonstrates that HSV-1 impacts promoter-proximal Pol II pausing in a more complex and unexpected manner than previously thought.

## Results

### Widespread changes in promoter-proximal Pol II pausing during HSV-1 infection

The standard measure for quantifying promoter-proximal pausing is the so-called pausing index (PI) of a gene, which is calculated as the ratio of normalized read counts in a window around the TSS (=promoter window) divided by normalized read counts in a window on the gene body excluding the promoter. PIs were also used by Rivas *et al*. to quantify the effects of lytic HSV-1 infection on Pol II pausing (9). We thus started by performing a genome-wide PI analysis using the published PRO-seq data of mock and 3 h p.i. WT-F infection (5). Since annotated gene 5’ends do not necessarily reflect the used TSS in a cell type and multiple alternative TSS are often annotated, we first identified the dominantly used TSS for each gene from published PROcap-seq and PRO-seq data of flavopiridol-treated uninfected human foreskin fibroblasts (HFF) (29) (see methods). PROcap-seq is a variation of PRO-seq that specifically maps Pol II initiation sites. Flavopiridol inhibits CDK9 and thus arrests Pol II in a paused state at the TSS (30) and allows also identifying the TSS for genes that are not or weakly paused in untreated cells. Consistent peaks in PROcap-seq and PRO-seq of flavopiridol-treated HFF provided an initial set of 136,090 putative TSS positions, which were further filtered to identify high confidence sites by requiring a maximum distance of 500 bp to the nearest annotated gene. This identified 42,193 potential TSS positions for 7,650 genes (median number of TSS per gene = 4 with a median distance of 42 bp). For each gene, the TSS with the highest expression was selected for further analyses. Although the PRO-seq data by Birkenheuer *et al*. was obtained in HEp-2 cells, for most genes the identified TSS in HFF matched very well to PRO-seq peaks in mock infected HEp-2 cells (**Fig. S1a** in the supplemental material), better than annotated gene 3’ends (**Fig. S1b**).

For PI calculation, normalized read counts were determined in a strand-specific manner as reads per kilobase million (RPKM) in the window from the TSS to TSS + 250 bp for the promoter region and from TSS + 250 bp to TSS +2,250 bp (or the gene 3’end if closer) for the gene body. It should be noted that there is no consensus on how to best define promoter and gene body windows for PI calculation and a wide range of alternative ranges have previously been used (see e.g., (31–33)). Genes with zero reads in the promoter or gene body window in mock or WT-F 3 h p.i. were excluded, resulting in PIs for 7,056 genes (**Data Set S1** in the supplemental material). This analysis showed that for the vast majority of genes PIs were reduced upon 3 h p.i. HSV-1 infection compared to mock (**Fig. 1a**). Even with very lenient criteria for an increase in PI, i.e., a fold-change > 1 in HSV-1 infection compared to mock, only 763 genes (10.8%) showed an increase in PI upon HSV-1 infection (red in **Fig. 1a**). In contrast, 2,082 genes (29.5%) exhibited a slightly reduced PI (fold-change < 1 but ≥ 0.5, blue in **Fig. 1a**) and 4,211 genes (59.7%) showed a strongly reduced PI (fold-change < 0.5, i.e. more than 2-fold reduced, green in **Fig. 1a**). Thus, HSV-1 infection induces widespread changes in promoter-proximal Pol II pausing of host genes, resulting in PI reductions for almost all genes and strong PI reductions for the majority of genes.

**Fig. 1:**
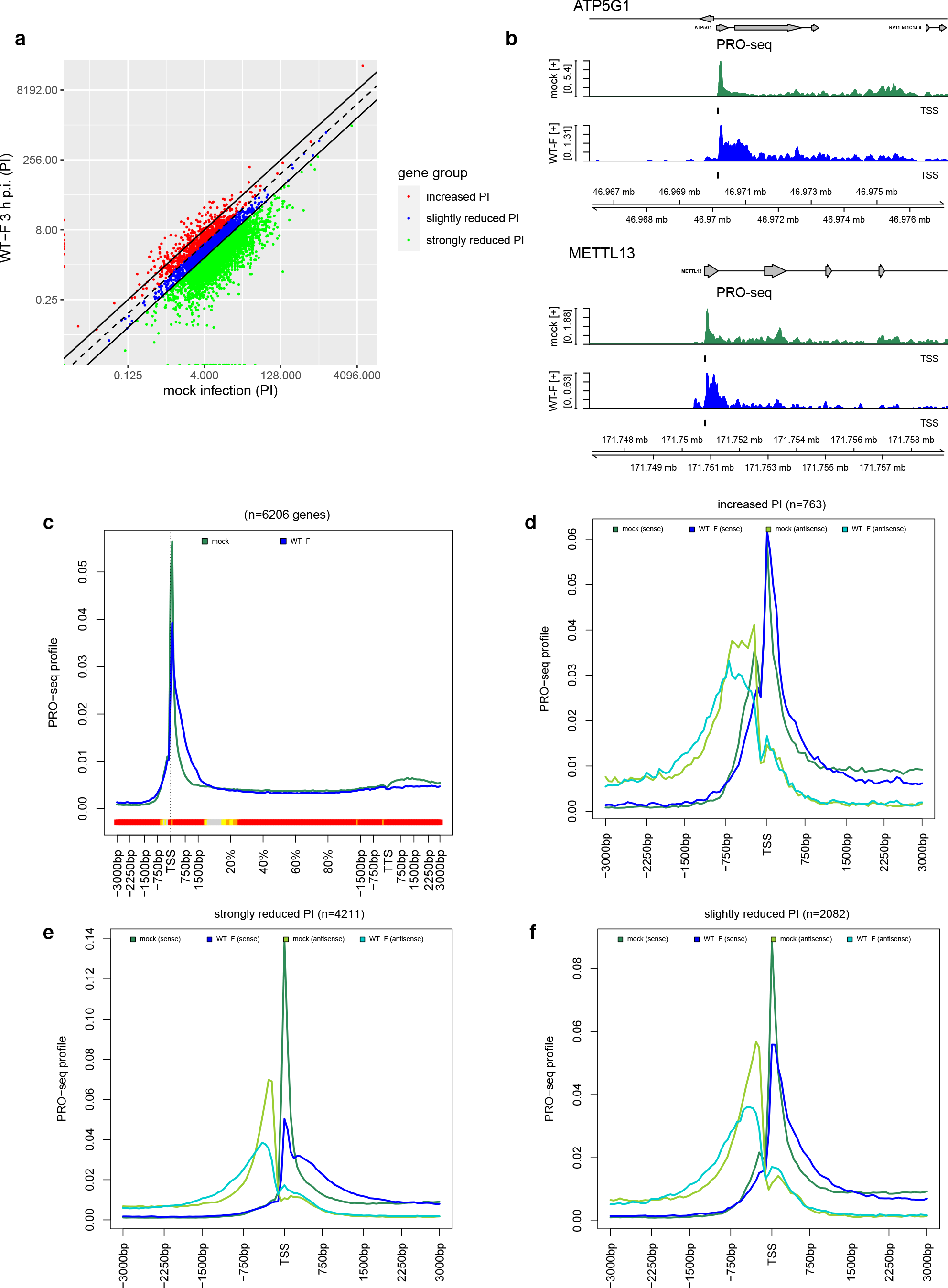
HSV-1 infection impacts promoter-proximal pausing of most host genes. **(a)** Scatter plots comparing pausing indices (PI) between mock and WT-F infection at 3 h p.i. The dashed line indicates equal PI values and solid lines a 2-fold change in PIs. Genes were divided into three groups according to changes in their PI: (1) increased PI in HSV-1 infection (fold-change >1, 763 genes, red), (2) slightly reduced PI in HSV-1 infection (fold-change < 1 but ≥ 0.5, 2,082 genes, blue), (3) strongly reduced PI in HSV-1 infection (fold-change < 0.5, 4,211 genes, green). **(b)** Read coverage around the TSS in PRO-Seq data (sense strand only) for mock (dark green) and WT-F infection at 3 h p.i. (dark blue) for example genes with a reduction in PI upon HSV-1 infection. Read coverage was normalized to total number of mapped reads and averaged between replicates. The identified TSS used in the analysis is indicated by a short vertical line below each read coverage track. Gene annotation is indicated at the top. Boxes represent exons, lines represent introns and direction is indicated by arrowheads. Genomic coordinates are shown on the bottom. Figures are not centered around the TSS, but a larger region downstream of the TSS was included than upstream of the TSS. **(c)** Metagene plot showing the distribution of PRO-seq profiles from 3 kb upstream of the TSS to 3 kb downstream of the TTS in sense direction for mock (dark green) and WT-F 3 h p.i. infection (dark blue) for all analyzed genes with a gene length > 3 kb. Here, regions from −3 kb to +1.5 kb of the TSS and from −1.5 kb to +3 kb of the TTS were divided into 90 bp bins, respectively, and the remainder of the gene body (+1.5 kb of TSS to −1.5 kb of TTS) into 100 bins of variable length to compare genes with different lengths. Shorter genes were excluded as regions around the TSS and TTS would overlap otherwise, resulting in 6,206 genes. The colored band below the metagene curves indicates the significance of paired Wilcoxon tests comparing the normalized PRO-seq coverages of genes for each bin between mock and WT-F 3 h p.i. infection. P□values are adjusted for multiple testing with the Bonferroni method within each subfigure; color code: red = adj. p-value ≤ 10^−15^, orange = adj. p-value ≤ 10^−10^, yellow = adj. p-value ≤ 10^−3^. **(d-f)** Metagene plots showing the distribution of PRO-seq profiles in sense (dark green and blue) and antisense (light green and blue) direction from −3 kb to + 3 kb around the TSS for the three gene groups defined in **(a)** with increased PI **(d)**, strongly reduced PI **(e)**, and slightly reduced PI **(f)**. Mock infection is shown in dark (sense) and light green (antisense) and WT-F infection at 3 h p.i. in dark (sense) and light blue (antisense). For this purpose, the TSS ± 3 kb promoter window for each gene was divided into 101 bins, PRO-seq read counts for each sample were determined for each bin, normalized to sequencing depth and averaged across replicates. Subsequently, bin values for each gene were normalized to sum up to 1 to obtain the Pol II occupancy profile in the promoter window. PRO-seq profiles were determined separately for sense and antisense strand.

### HSV-1 infection shifts Pol II pausing to downstream sites for most host genes

A disadvantage of PI analysis is that PIs are not only impacted by increases or decreases in Pol II pausing with decreased or increased elongation across the gene body, but by any alteration in Pol II occupancy affecting the number of reads in either promoter or gene body windows. We thus next investigated read coverage in a genome viewer for several example genes with reduced PI. This indicated that changes in promoter-proximal Pol II pausing during HSV-1 infection were highly complex as exemplified by the ATP5G1 and METTL13 genes in **Fig. 1b**. Instead of narrow promoter-proximal PRO-seq peaks observed in mock infection, promoter peaks in HSV-1 infection often extended into the gene body by a few hundred nucleotides (e.g., ATP5G1) and/or additional downstream peaks were observed as e.g., for METTL13. However, at the end of these extended/additional peaks read levels dropped again to similarly low levels relative to the promoter peak as in uninfected cells. Read levels were not increased across the whole gene body relative to the promoter peak as would be expected with increased levels of elongation and productive transcription. The extended promoter peaks and additional downstream peaks in HSV-1 infection reduced PI values, as they extended >250 bp from the TSS into the gene body. Consequently, gene body RPKM, i.e., the denominator in PI calculation, was increased relative to the promotor RPKM, i.e., the numerator. It should be noted that overall Pol II occupancy was reduced both on the promoter and gene body for the majority of genes during HSV-1 infection as already reported by Birkenheuer *et al*. (5). To allow comparing the distribution of Pol II occupancy, not absolute levels, in read coverage plots like **Fig. 1b** and visualize downstream shifts in pausing sites, different scales are used for mock and HSV-1 infection. These reflect the highest values observed in mock and HSV-1 infection, respectively, for the selected genomic region.

To investigate whether such complex pausing changes were a global trend, we performed metagene analyses in ± 3 kb windows around promoters for all 7,650 analyzed genes (**Fig. S2**, excluding 1 gene without reads in any sample) as well as separately for the three gene groups defined above based on PI changes (**Fig. 1d-f**). For metagene analyses, the 6 kb promoter windows for each gene were divided into 101 bins. PRO-seq read counts were determined for each bin in a strand-specific manner, normalized to sequencing depth and averaged across replicates. Subsequently, bin values for each gene were normalized to sum up to 1 to obtain the Pol II occupancy profile around the promoter for each gene before averaging across all genes. This normalization allows comparing Pol II occupancy around the promoter between genes with different expression levels and makes the analysis independent of global changes in Pol II occupancy between mock and HSV-1 infection. As a consequence, sharp, singular peaks at promoters are characterized by higher peak maxima (e.g., dark green curves in **Fig. 1c**), while broader peaks or multiple peaks have lower peak maxima (e.g., dark blue curves in **Fig. 1c,e**). Normalization was performed independently for sense and antisense PRO-seq profiles, thus the height of peaks does not reflect relative levels of sense vs. antisense transcription but the distribution of sense and antisense transcription around the TSS, respectively. Analysis of all genes already showed that PRO-seq profiles for both sense and antisense direction were significantly altered between mock and HSV-1 infection (**Fig. S2**). In HSV-1 infection, lower Pol II occupancy was observed directly at the TSS and increased occupancy down- and upstream of the TSS for sense and antisense transcription, respectively. This was limited to within 2,250 bp of the TSS in both cases. Metagene analysis on complete genes from the promoter to downstream of the transcription termination site (TTS) confirmed that this relative increase in occupancy downstream of the TSS did not extend across gene bodies (**Fig. 1c**). To assess significance of differences between two conditions, Wilcoxon signed rank tests were performed for each bin in metagene plots comparing normalized coverage values for each gene between the two conditions across all genes. Multiple testing corrected p-values are color-coded at the bottom of metagene plots (red = adj. p-value ≤ 10^−15^, orange = adj. p-value ≤ 10^−10^, yellow = adj. p-value ≤ 10^−3^, p-values not shown if >2 curves are included in metagene plots). This showed highly significant differences in the distribution of Pol II occupancy between mock and WT infection for almost the complete gene body, with the notable exception of the region at the end of the promoter window, where increased relative Pol II occupancy in HSV-1 infection downstream of the TSS changed to decreased relative Pol II occupancy on the gene body. The reduction in Pol II occupancy downstream of the TTS during HSV-1 infection reflects loss of Pol II pausing at the TTS associated with disruption of transcription termination previously reported in HSV-1 infection (8). Interestingly, genes with increased PI upon HSV-1 infection (red in **Fig. 1a**) only showed a small change in Pol II occupancy in sense direction at the promoter in HSV-1 infection (**Fig. 1d**). In contrast, genes with strong PI reduction upon HSV-1 infection showed a strong reduction of the major peak height at the TSS, a pronounced broadening of the peak into the gene body and a second minor peak downstream the TSS (**Fig. 1e**). A similar but less pronounced effect was observed for genes with a weak reduction in PI, with a general broadening of the TSS peak but no minor peak (**Fig. 1f**). These changes in the distribution of Pol II occupancy explain the reduction in PIs as read counts in the in the gene body window are increased relative to read counts in the promoter window.

To identify groups of genes with distinct patterns of changes of Pol II occupancy around the TSS between HSV-1 and mock infection, we performed hierarchical clustering of genes based on their PRO-seq profiles in sense direction for both mock and HSV-1 infection (**Fig. 2a**). Since we wanted to ensure that genes with distinct patterns of changes were placed in different clusters, a stringent cutoff was applied on the clustering dendrogram to obtain 50 gene clusters at the cost of obtaining multiple clusters with similar patterns. While most clusters exhibited only a narrow peak at the TSS in mock infection (e.g., **Fig. 2b, Fig. S3a,b,f-h**), some already exhibited a second minor peak shortly after the TSS already prior to infection (e.g., **Fig. 2c,d, S3c-e**). In addition, a few clusters representing a total of 2,018 genes showed peaks in mock infection that were shifted relative to the TSS we had identified (e.g., **Fig. 2d**, **Fig. S3e**). In all cases, the peak was shifted at most 750 bp from the identified TSS and was commonly within 100-200 bp of the identified TSS. Since position of peaks within clusters were highly similar due to the stringent clustering cutoff, analysis of individual clusters thus avoids confounding effects resulting from misidentification of TSS positions. Typical Pol II occupancy changes upon HSV-1 infection included a reduction of peak height at the TSS with a broadening of the peak into the gene body (e.g., **Fig. 2b**), changes in minor and major peak heights in case multiple peaks were already present prior to infection (e.g., **Fig. 2c,d**) as well as new peaks originating downstream of the TSS in HSV-1 infection (e.g., **Fig. 2e**). **Fig. S4** provides an overview on positions, number and relative heights of Pol II occupancy peaks in mock and HSV-1 infection for each cluster. In total, 30 clusters shared the same major TSS peak between mock and HSV-1 infection, which included also clusters with only a reduction in peak height but no broadening of the TSS peak (Cluster 20; **Fig. S3f**) and clusters without loss of Pol II pausing (Clusters 21 & 27; **Fig. S3g,h**). 28 clusters showed a second peak in HSV-1 infection downstream of the TSS peak with a median distance of 480 bp to the major peak (e.g., **Fig. 2c,d**). For 9 of these 28 clusters, the major TSS peak differed between mock and HSV-1 infection (e.g., **Fig. 2e**). Almost all clusters exhibited a reduced Pol II peak height at the TSS (except Clusters 27 & 21, 262 genes; **Fig. S3g,h**) and most clusters with a reduced Pol II peak height showed an extension of the peak into the gene body or an increased downstream peak (except Clusters 7, 20, 23, 24 & 36, 480 genes). Read coverage plots for example genes from different clusters are shown in **Fig. S5** and a UCSC browser session showing PRO-seq read coverage for all human genes separately for replicates is available at https://genome.ucsc.edu/s/Caroline%20Friedel/PROseq_HSV1.

**Fig. 2:**
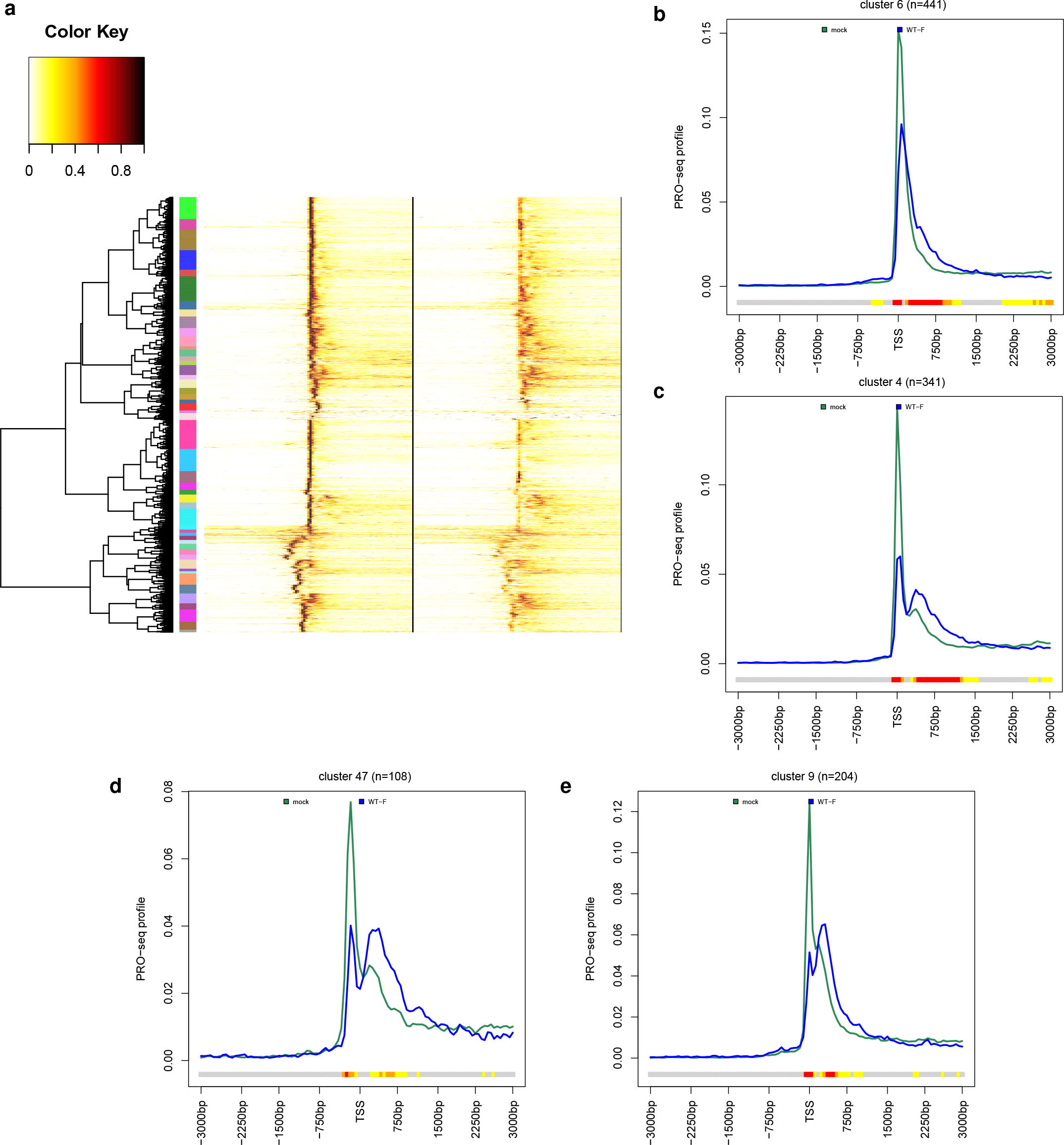
Distinct patterns of changes in promoter-proximal pausing upon HSV-1 infection. **(a)** Heatmap showing the result of the hierarchical clustering analysis of PRO-seq profiles in mock and WT-F infection. For clustering, PRO-seq profiles in sense direction for mock and WT-F infection were first concatenated and then divided by the maximum value in the concatenated profiles. This resulted in a value of 1 for the position of the highest peak in either mock or HSV-1 infection. Hierarchical clustering was performed according to Euclidean distances and Ward’s clustering criterion and the cutoff on the hierarchical clustering dendrogram was selected to obtain 50 clusters (marked by colored rectangles between the dendrogram and heatmap). Clusters are numbered from top to bottom. **(b-e)** Metagene plots of PRO-seq profiles on the sense strand in mock (dark green) and WT-F infection at 3 h p.i. (dark blue) for example Clusters 6, 4, 47 and 9 from **(a)**. See methods and legend to **Fig. 1** for an explanation of metagene plots. The colored bands below the metagene curves in each panel indicate the significance of paired Wilcoxon tests comparing the normalized PRO-seq coverages of genes for each bin between mock and WT-F 3 h p.i. infection. P□values are adjusted for multiple testing with the Bonferroni method within each subfigure; color code: red = adj. p-value ≤ 10^−15^, orange = adj. p-value ≤ 10^−10^, yellow = adj. p-value ≤ 10^−3^.

To investigate whether the different patterns in Pol II pausing changes were correlated to gene function or transcription factor binding, we performed over- and under-representation analysis for Gene Ontology (GO) terms and transcription factor binding motifs from TRANSFAC for each cluster (Data Set S2, adj. p-value cutoff < 0.001, a stringent p-value cutoff was chosen to adjust for performing this analysis separately for 50 clusters). This revealed an enrichment of subunits of the spliceosomal snRNP complex, specifically U5 and U6 snRNAs, in Cluster 27 (adj. p-value < 1.31 × 10^−7^), one of the clusters without change in pausing. Furthermore, Cluster 16 was enriched for genes encoding proteins of the large ribosomal subunit (adj. p-value < 0.00057). For both Clusters, < 8.5% of genes belonged to the enriched GO categories and no other over- or under-representation was observed. Thus, clusters identified based on changes in pausing did not represent functionally related gene groups. Interestingly, however, Cluster 32 was strongly enriched for a number of G- or C-rich transcription factor binding motifs, with 90% of genes having a match for a long G-rich motif (GGGMGGGGSSGGGGGGGGGGGG, adj. p-value <0.00025). In contrast, several A-and T-rich motifs (e.g. NNNNRNTAATTARY, adj. p-value < 6.94 × 10^−9^) were under-represented in Cluster 32. The opposite effect was observed for Cluster 6, with G-/C-rich motifs being under- and A-/T-rich motifs being over-represented. A few G-/C-rich motifs were also under-represented in Cluster 10. Recently, Watts *et al*. showed that GC content is high around pause sites and that GC skew (= (G-C)/(G+C)) peaks in the 100nt upstream of the pause site (34). Analysis of GC content and GC skew around the TSS for individual clusters indeed showed a high GC content for Cluster 32 at and downstream of the TSS, though no peak in GC skew (**Fig. S6a**). In contrast, GC content was less increased around the TSS for Cluster 6, while GC skew peaked at the TSS and was increased downstream of the TSS (**Fig. S6b**). As Clusters 32 and 6 differed considerably regarding the change in Pol II pausing upon HSV-1 infection, this raises the possibility that sequence composition around the TSS could play a role in determining the changes in Pol II pausing upon HSV-1 infection. However, analysis of the other clusters did not reveal any consistent trend in GC content or GC skew that explained differences in Pol II pausing between clusters. The one consistent trend we observed was that clusters for which the major PRO-seq peak was significantly upstream of our identified TSS commonly exhibited a plateau of high GC content starting at the PRO-seq peak and extending to the TSS (e.g. **Fig. S6c,d**).

Considering the bidirectionality of transcription initiation at human promoters, we also performed metagene analyses of PRO-seq profiles in antisense direction. Antisense transcription initiation at bidirectional promoters commonly only results in short, unspliced, non-polyadenylated and unstable upstream antisense RNAs (uaRNAs) (35), that have highly heterogeneous 3′ ends (36). The metagene analysis on all genes of antisense PRO-seq profiles also showed a reduction in the antisense TSS peak height and a broadening of the peak in antisense direction (**Fig. S2**). However, clustering of antisense PRO-seq profiles in mock and HSV-1 infection with the same approach as for sense profiles to obtain 50 clusters did not identify different patterns between clusters. Most of the 50 antisense clusters exhibited only the same pattern as the metagene analysis of all genes (e.g., **Fig. 3a**). Only two clusters (430 & 375 genes) showed a small secondary antisense peak originating in HSV-1 infection (**Fig. 3b,c**), while another cluster (129 genes) showed a secondary peak that was already present in mock infection but increased relative to the TSS peak in WT infection (**Fig. 3d**).

**Fig. 3:**
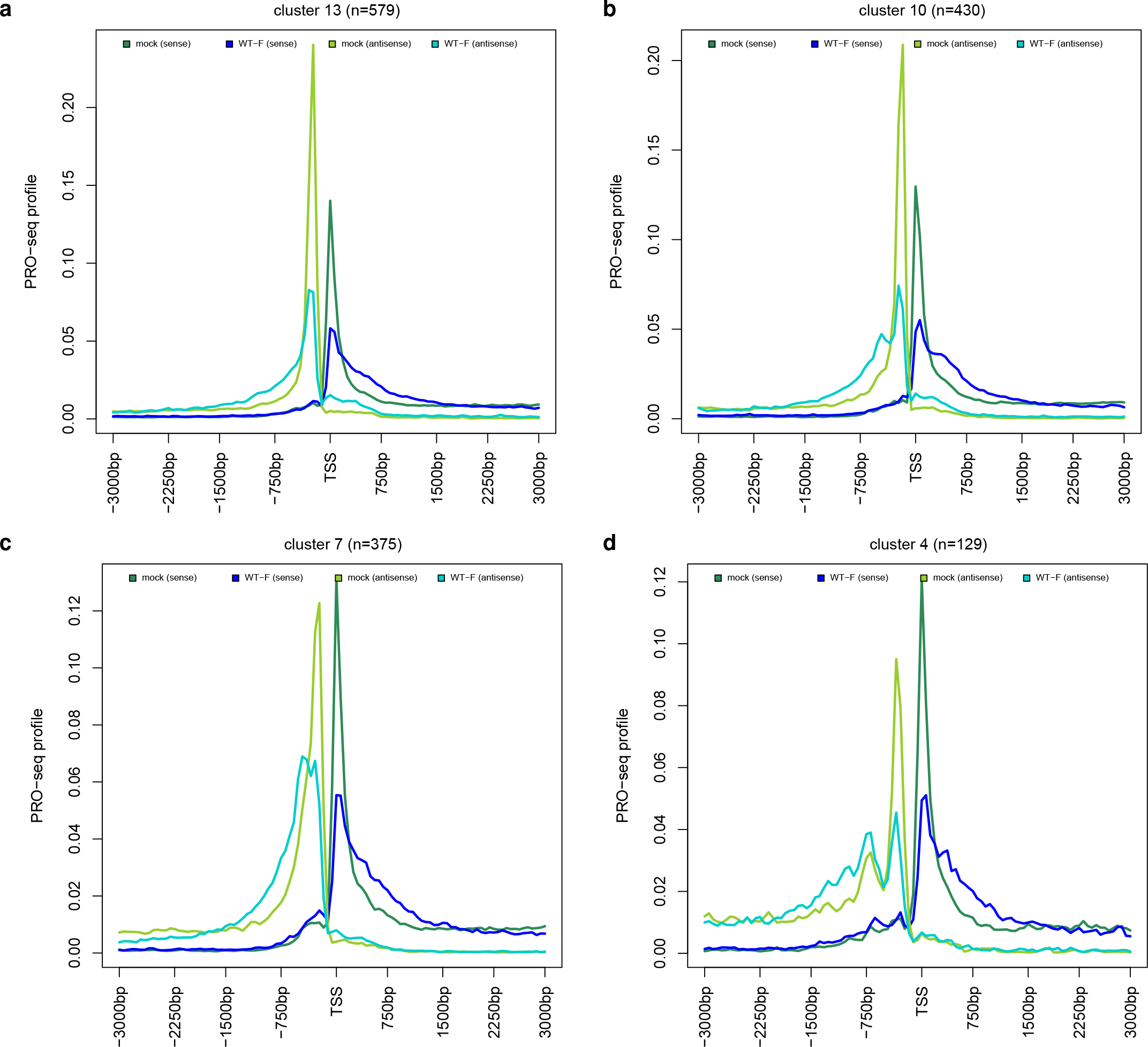
Changes in antisense promoter-proximal pausing in HSV-1 infection. Metagene plots of PRO-seq profiles on both sense (dark green and blue) and antisense (light green and blue) direction in mock (dark and light green) and WT-F infection at 3 h p.i. (dark and light blue) for example clusters resulting from the hierarchical clustering of genes according to antisense PRO-seq profiles in mock and WT-F infection. Here, clustering was performed as described in the legend for **Fig. 2a** but applied to concatenated antisense PRO-seq profiles in mock and WT-F infection. Thus, clusters shown here differ from the clusters shown in all other figures. **(a)** shows the most common pattern observed for almost all clusters with a broadening of the antisense PRO-seq peak at the TSS, while **(b-c)** show the only three clusters that exhibit different patterns with additional peaks originating or increasing in antisense direction during infection.

### Delayed pausing is not an artifact of *de novo* transcription initiation or read-through transcription

Since increasing or newly originating secondary PRO-seq peaks could also represent alternative transcription initiation, we next investigated the presence of alternative TSSs for all clusters in either the PROcap-seq and PRO-seq data of flavopiridol-treated HFF or the human genome annotation. For most clusters, <15% of genes showed evidence for an alternative TSS at the additional peak positions in either flavopiridol-treated cells (e.g., **Fig. S7a,c**) or the genome annotation (e.g., **Fig. S7b,d**). This is in clear contrast to clusters in which the TSS identified from flavopiridol-treated HFF did not represent the dominant TSS in HEp-2 cells. Here, almost 50% of genes had an additional peak in flavopiridol-treated cells or an annotated transcript start at the position of the dominant TSS in HEp-2 cells (e.g., **Fig. S7e,f**). We furthermore investigated induction of alternative *de novo* transcription initiation downstream of the TSS during HSV-1 infection using dRNA- and cRNA-seq data of transcript 5’ends for mock and HSV-1 strain 17 (WT-17) infection of HFF from our recent re-annotation of the HSV-1 genome (n=2 replicates) (37). dRNA-seq is based on selective cloning and sequencing of the 5′-ends of cap-protected RNA molecules resistant to the 5′–3′-exonuclease XRN1. cRNA-seq is based on circularization of RNA fragments. Both methods strongly enrich reads from 5′ RNA ends. dRNA-seq was performed for mock and 8 h p.i. HSV-1 infection with and without XRN1 treatment. cRNA-seq was performed for mock, 1, 2, 4, 6 and 8 h p.i. HSV-1 infection. Metagene analyses of dRNA- and cRNA-seq data showed clear peaks coinciding with the major PRO-seq peaks in mock infection and smaller peaks at minor PRO-seq peaks already present in mock infection (**Fig. 4a-d**, **Fig. S8, S9**). In contrast, no (increased) peaks were observed at the positions of downstream PRO-seq peaks that increased or newly originated during HSV-1 infection. In summary, changes in Pol II occupancy during HSV-1 infection are not due to alternative initiation at novel TSSs leading to capped transcripts, however we cannot completely exclude that they may reflect abortive *de novo* initiation at novel initiation sites downstream of the TSS. This analysis also excludes Pol II creeping, which is observed upon H2O2 treatment (38), as the latter would lead to signals from capped transcripts increasing downstream of the TSS in the pausing region.

**Fig. 4:**
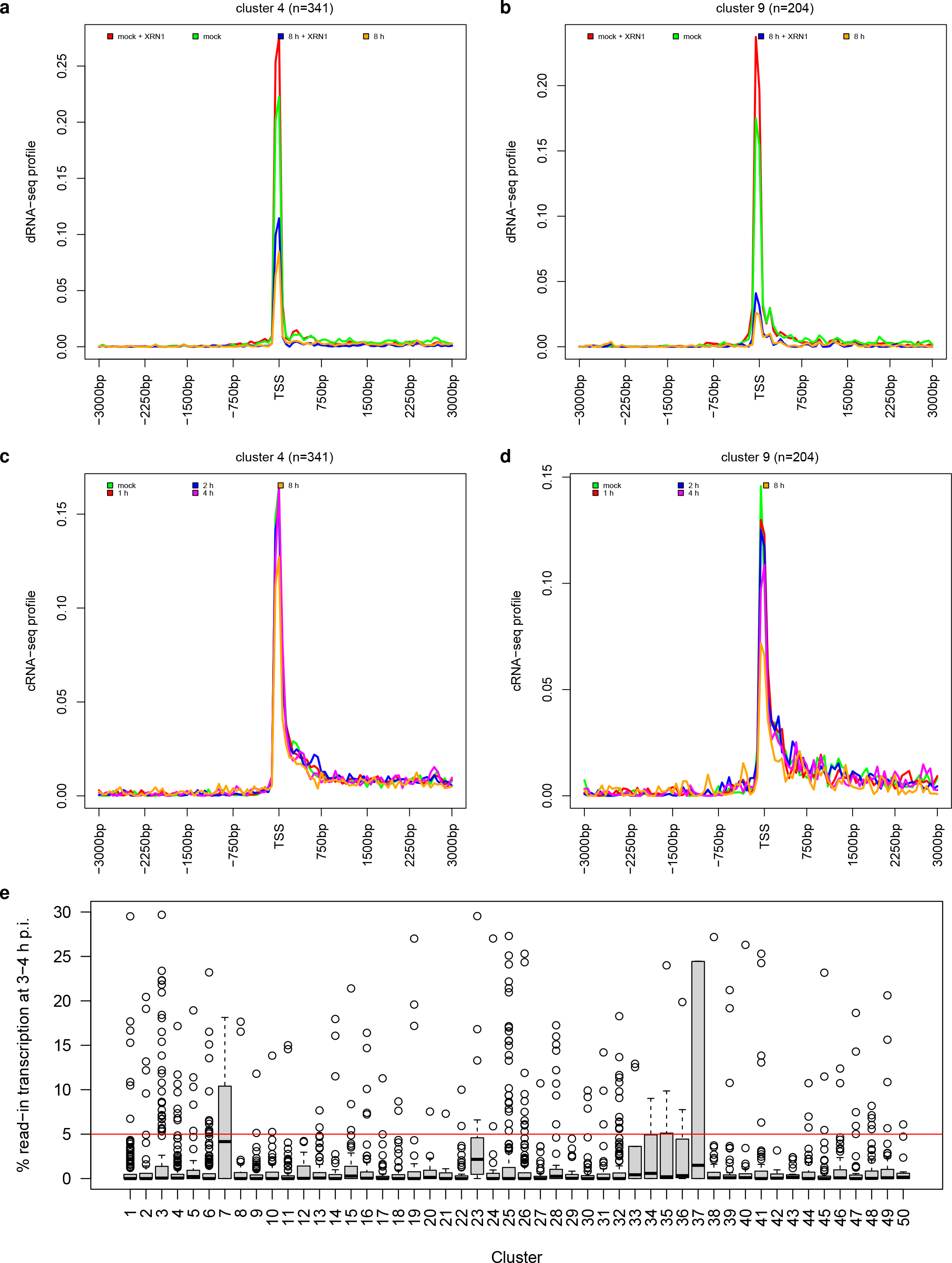
Delayed pausing is not an artefact of alternative *de novo* initiation or read-in transcription. **(a-d)** Metagene plots of **(a,b)** dRNA-seq profiles on the sense strand in mock and WT-17 8 h p.i. infection with and without XRN1 treatment and **(c,d)** cRNA-seq profiles on the sense strand in mock and WT-17 infection at 1, 2, 4, and 8 h p.i. for example Clusters 4 and 9, which show broadening of peaks or additional peaks originating or increasing in height in PRO-seq data during WT-F infection. For metagene plots of PRO-seq profiles for these clusters see **Fig. 2c,d**. **(e)** Boxplots showing the distribution of read-in transcription at 3-4 h p.i. for genes in the 50 clusters identified from sense PRO-seq profiles. Boxes represent the range between the first and third quartiles for each cluster. Black horizontal lines in boxes show the median. The ends of the whiskers (vertical lines) extend the box by 1.5 times the inter-quartile range. Data points outside this range (outliers) are shown as small circles. The red horizontal line indicates the cutoff we previously used to determine that no read-in transcription is observed (≤5% read-in transcription). Metagene plots of PRO-seq profiles for clusters 7, 23 and 33-37 with some read-in transcription observed at 3-4 h p.i. are shown in **Fig. S10**.

We previously showed that late in infection “read-in” transcription originating from disrupted transcription termination for an upstream gene can extend into downstream genes, which can be mistaken for induction of the downstream gene (8). Although read-in transcription only affected very few genes within the first 4 h p.i., increased Pol II occupancy downstream of the TSS could potentially also originate from read-in transcription. To quantify read-in transcription, we used our previously published 4sU-seq time-course for every hour of the first 8 h of lytic HSV-1 strain 17 (WT-17) infection of HFF (8). 4sU-seq sequences newly transcribed RNA obtained by labeling with 4-thiouridine (4sU) in specific time intervals of infection (here: 1 h intervals for the first 8 h of lytic expression). Read-in transcription was quantified as previously described (see (7, 39) and Materials and Methods for details). In brief, we first calculated the percentage of upstream transcription (= transcription in a 5 kb window upstream of the gene 3’end / gene expression) for mock infection and each 1 h window of HSV-1 infection. Subsequently, percentage of read-in transcription was calculated by subtracting values in mock infection from values in each 1 h window of HSV-1 infection (multiplied by 100, negative values set to zero). This analysis included only genes with ≥ 5 kb to the next up- or downstream gene. By 3-4 h p.i., read-in transcription was essentially absent (i.e., much less than 5%) for almost all genes in almost all clusters (**Fig. 4e**). Only few clusters (Clusters 7, 23, 33-37) exhibited a small extent of read-in transcription already this early in infection, however, these clusters did not exhibit substantial downstream shifts in Pol II occupancy or additional secondary peaks (**Fig. S10**). The largest of these clusters, Cluster 7, indeed showed significantly increased Pol II occupancy in sense direction upstream of the TSS in HSV-1 infection (**Fig. S10a**), consistent with read-in transcription. We conclude that read-in transcription extending (partially) into downstream genes does not explain extended TSS peaks or novel or increasing downstream peaks in Pol II occupancy observed in HSV-1 infection.

### Delayed pausing in HSV-1 infection occurs downstream of 2^nd^ pause sites used upon NELF depletion

Recently, Aoi *et al*. showed that rapid depletion of NELF, the key mediator of Pol II pausing, does not completely abolish pausing (28). Instead, Pol II is paused at a 2^nd^ more downstream pause site around the +1 nucleosome. Since Rivas *et al*. showed an ICP4-dependent decrease of NELF in the promoter-proximal region of some HSV-1-activated genes (9), we reanalyzed PRO-seq data from the Aoi *et al*. study for 0, 1, 2 and 4 h auxin-induced degradation of NELF for our 50 clusters to investigate whether changes in pausing upon NELF depletion showed similarities to changes of HSV-1 infection. For a few of our clusters, metagene analyses indeed showed an increased second PRO-seq peak upon NELF degradation or a broadening of the first PRO-seq peak downstream of the TSS (e.g., **Fig. 5, Fig. S11a-d**). For most clusters, however, we only observed a reduction in the major peak height and a minor broadening of the peak into downstream regions (e.g., **Fig. S11e-j**). In either case, the changes in the distribution of Pol II occupancy in HSV-1 infection were much more pronounced than after NELF depletion, with more extensive broadening of peaks and new secondary peaks being further downstream of the major peak. In summary, delayed pause sites in HSV-1 infection are further downstream than “normal” 2^nd^ pause sites at +1 nucleosome positions used upon NELF depletion.

**Figure. 5:**
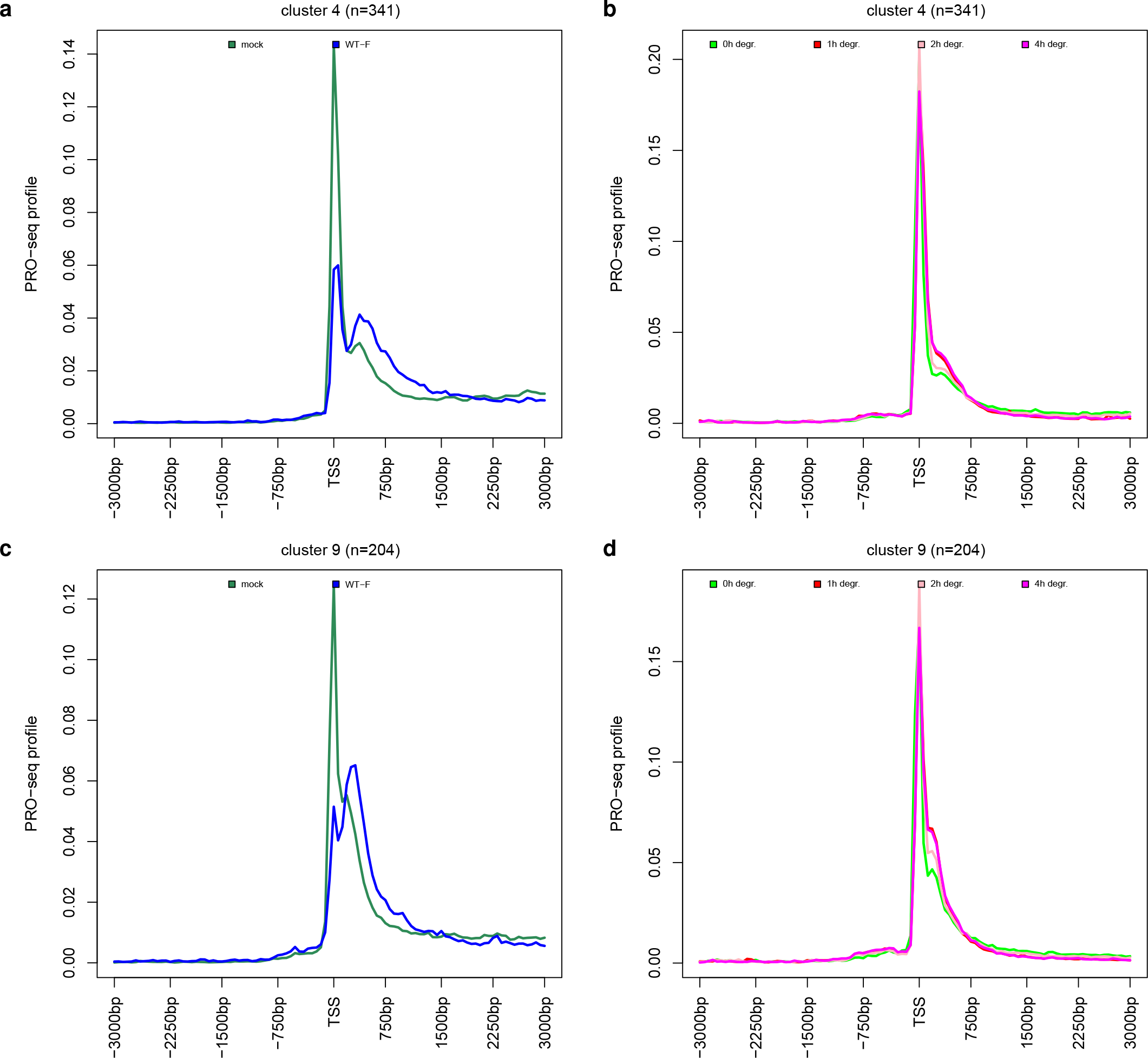
HSV-1 infection leads to stronger downstream shifts in pause sites than NELF depletion. Metagene plots around the TSS of PRO-Seq profiles for **(a,c)** mock and WT-F 3 h p.i. infection from the study of Birkenheuer *et al*. (5) and **(b,d)** 0, 1, 2, and 4 h auxin-inducible degradation of NELF from the study by Aoi *et al*. (28) for example clusters showing a broadening of the TSS peak **(a,b)** or a small downstream peak **(c,d)** upon NELF degradation.

## Discussion

Promoter-proximal Pol II pausing is a key regulatory step between transcription initiation and productive elongation. HSV-1 infection has previously been reported to dramatically impact Pol II positioning on host genes, including promoter-proximal regions (5). Promoter-proximal Pol II pausing on HSV-1 genes also plays a key role in HSV-1 transcription (27). Rivas *et al*. recently reported that HSV-1 infection leads to a reduction in pausing indices for activated host genes and an increase in pausing indices at repressed host genes (9). While our re-analysis of PRO-seq data of mock and WT-F 3 h p.i. infection also showed a reduction of pausing indices for most expressed host genes during HSV-1 infection, it also illustrated that pausing indices are an imperfect though easy-to-use measure of promoter-proximal Pol II pausing. Pausing indices are altered by any change in the distribution of Pol II between promoter and gene body. Thus, more in-depth analyses are necessary to characterize changes in promoter-proximal Pol II pausing, not only during HSV-1 infection. Our metagene analyses revealed that HSV-1 infection does not lead to a simple reduction of promoter-proximal Pol II pausing with a relative increase of elongating Pol II on the whole gene body. Instead, we observed that Pol II pausing is retained for the vast majority of genes but is shifted to downstream pause sites. This is reflected in broadened Pol II promoter-proximal peaks that extend further into the gene body and newly originating or increasing downstream peaks. A fine-grained clustering analysis identified a wide range of different patterns for different genes in HSV-1 infection, which contrasts with the sharp promoter-proximal peaks commonly observed in uninfected cells. This indicates that positioning of shifted pause sites in HSV-1 infection is less well-defined than of “normal” pause sites in uninfected cells. Analysis of transcript start site profiling and newly transcribed RNA in HSV-1 infection excluded that this was due to *de novo* initiation at downstream sites or read-in transcription originating from disrupted transcription termination for upstream genes.

Interestingly, analysis of promoter-proximal pausing in antisense direction also showed a broadening of antisense TSS peaks upon HSV-1 infection, with antisense transcription extending further upstream of the TSS than in uninfected cells. However, secondary antisense peaks were only observed for a small fraction of genes (12.2%). It has been proposed that Pol II is particularly prone to pausing and termination during early elongation, specifically on AT-rich sequences often found upstream of promoters (40). Previously, we reported both widespread disruption of transcription termination in HSV-1 infection (8) and activation of antisense transcription at promoters and within gene bodies (41). It is thus tempting to speculate that activation of antisense transcription in HSV-1 infection could be linked to alterations in antisense Pol II pausing, potentially in combination with disruption of transcription termination in antisense direction.

Nucleosomes represent a natural barrier to transcription and are disassembled before and reassembled after transcribing Pol II (42). Nucleosomes directly downstream of the TSS are generally well-positioned at specific locations, in particular the +1 nucleosome, but less so further up- or downstream (43–45). In presence of NELF, Pol II pausing occurs between the promoter and the +1 nucleosome, and strong positioning of the +1 nucleosome increases pausing (46). While NELF has previously been considered to be required for establishing Pol II pausing, rapid depletion of NELF using auxin-inducible degron does not abolish pausing (28). Instead, pausing appears to be a two-step process with Pol II transitioning from the 1^st^ to a 2^nd^ pause site associated with +1 nucleosomes upon NELF depletion. As Rivas *et al*. reported decreased levels of NELF at promoter regions of four activated genes tested by ChIP, depletion of NELF from host promoters could be implicated in the downstream shift of promoter-proximal Pol II pausing in HSV-1 infection. Notably, a few clusters with additional downstream peaks observed in HSV-1 infection already showed small and much less pronounced peaks at these positions in mock infection. This puts some weight to the hypothesis that loss of pausing at major pause sites upon HSV-1 infection leads to use of 2^nd^ downstream pause sites. However, comparison of the effects of NELF degradation and HSV-1 infection for the 50 identified clusters showed that HSV-1 infection led to much more pronounced alterations in Pol II pausing and more extensive downstream shifts of pause sites than degradation of NELF. Thus, NELF depletion at promoters alone cannot explain the delay in Pol II pausing observed in HSV-1 infection, though it may contribute.

The role of ICP22 in shaping Pol II pausing during WT HSV-1 infection also remains an open and intriguing question. ICP22 directly interacts with the P-TEFb subunit CDK9 to inhibit Pol II transcription elongation on host genes (20) and ectopic expression of an ICP22 segment interacting with CDK9 increases Pol II pausing similar to CDK9 inhibition (21). In the context of wild-type HSV-1 infection, however, this ICP22 activity appears insufficient to increase pausing for the majority of host genes. It could, however, contribute to establishment of Pol II pausing at downstream pause sites by hindering a CDK9-dependent switch to active elongation.

The functional role of the observed changes in promoter-proximal Pol II pausing during HSV-1 infection also remains unclear. Rivas *et al*. concluded that ICP4 activates host genes by promoting release of paused Pol II into elongation. However, our analysis showed that Pol II is not fully released from pausing but pausing is shifted to downstream sites for most genes. Thus, alterations in Pol II pausing do not generally lead to increased elongation and are furthermore not limited to activated genes. While this could still serve to promote increased elongation for a few genes, it could also simply be a by-product of other processes ongoing in HSV-1 infection, e.g., the general loss of Pol II from the host genome and its recruitment to viral genomes. Finally, it should be noted that our study has also important implications for analysis of functional genomics studies of HSV-1 and potentially other viral infections. As already observed in previous studies reporting on disruption of transcription termination or activation of antisense transcription observed upon HSV-1 infection (8, 41), standard sequencing data analysis methods are not designed and thus insufficient to uncover previously unsuspected alterations in transcription. Thus, more in-depth analyses and customized methods are required. In summary, our study highlights a novel aspect in which HSV-1 infection fundamentally alters the host transcriptional cycle, which has implications for our understanding not only of HSV-1 infection but also of maintenance of Pol II pausing in eukaryotic cells.

## Materials and Methods

### Previously published sequencing data analyzed in this study

PROcap-seq and PRO-seq data of flavopiridol-treated uninfected HFF cells were taken from the study by Parida *et al*. (29) (GEO accession: GSE113394, samples GSM3104917 and GSM3104913). PRO-seq data for mock and WT-F infection at 3 h p.i. of HEp-2 cells were taken from the study by Birkenheuer *et al*. (5) (n=3 replicates, GEO accession: GSE106126, samples GSM2830123 - GSM2830127). PRO-seq data for 0, 1, 2 and 4 h auxin-induced degradation of NELF were taken from the study by Aoi *et al*. (28) (n=1 apart from 0 h with n=2, GEO accession: GSE144786, samples GSM4296314 - GSM4296316, GSM4296318, GSM4296319). dRNA-seq data for mock and 8 h p.i. HSV-1 infection with and without XRN1 treatment and and cRNA-seq data for mock, 1, 2, 4, 6 and 8 h p.i. HSV-1 infection of HFFF were taken from our previous study (37) (n=2, GSE128324, samples GSM3671394 - GSM3671411). 4sU-seq data for mock and hourly intervals for the first 8 h of WT-17 infection of HFFF was taken from our previous study (8) (n=2, GEO accession: GSE59717).

### Read alignment

The read alignment pipeline was implemented and run in the workflow management system Watchdog (47, 48). Published sequencing data were first downloaded from SRA using the sratoolkit version 2.10.8. Sequencing reads were aligned against the human genome (GRCh37/hg19) and human rRNA sequences using ContextMap2 version 2.7.9 (49) (using BWA as short read aligner (50) and allowing a maximum indel size of 3 and at most 5 mismatches). For sequencing data of HSV-1 infection, alignment also included the HSV-1 genome (Human herpesvirus 1 strain 17, GenBank accession code: JN555585). For the two repeat regions in the HSV-1 genome, only one copy was retained each, excluding nucleotides 1–9,213 and 145,590–152,222 from the alignment. SAM output files of ContextMap2 were converted to BAM files using samtools (51). Read coverage in bedGraph format was calculated from BAM files using BEDTools (52).

### Data plotting and statistical analysis

All figures were created in R and all statistical analyses were performed in R (53). Read coverage plots were created using the R Bioconductor package Gviz (54).

### Transcription start site identification

We used the iTiSS program to identify candidate TSS in PROcap-seq and PRO-seq of flavopiridol-treated HFF (37, 55). For this purpose, iTiSS was run separately for each sample in the SPARSE_PEAK mode with standard parameters. Afterwards, the iTiSS TSRMerger program was used to select only peaks that were identified in both samples within +/− 5 bp. Consistent peaks were only further considered if they were within 500 bp to the nearest annotated gene and for each gene the TSS with the highest read count (weighted by the number of possible alignments for the read) was selected for further analyses.

### Calculation of pausing indices

PRO-seq read counts in promoter windows (TSS to TSS + 250 bp) and gene bodies (TSS + 250 bp to TSS + 2,250 bp or gene 3’ end if closer) were determined using featureCounts (56) and gene annotations from Ensembl (version 87 for GRCh37) (57) in a strand-specific manner and normalized by the total number of reads and window lengths to obtain RPKM values. RPKM values were averaged between replicates and genes with zero reads in either promoter or gene body window were excluded from analysis. PI for a gene was then calculated as the ratio of promoter RPKM to gene body RPKM.

### Metagene and clustering analysis

Metagene analyses were performed as previously described (58) using the R program developed for this previous publication (available with the Watchdog *binGenome* module in the Watchdog module repository (https://github.com/watchdog-wms/watchdog-wms-modules/)). For promoter region analyses, the regions −3 kb to +3 kb of the TSS were divided into 101 bp bins for each gene. For each bin, the average coverage per genome position was calculated in a strand-specific manner for PRO-seq data and bin read coverages were then normalized by dividing by the total sum of all bins. Metagene curves for each replicate were created by averaging results for corresponding bins across all genes and metagene plots then show the average metagene curves across replicates. Genes without any reads in any of the analyzed samples were excluded from the analysis. For metagene analyses on the whole gene, the regions from −3 kb to +1.5 kb of the TSS and from −1.5 kb to +3 kb of the TTS were divided into 90 bp bins and the remainder of the gene body (+1.5 kb of TSS to −1.5 kb of TTS) into 100 bins of variable length in order to compare genes with different lengths. Genes with a gene length < 3 kb were excluded as regions around the TSS and TTS would overlap otherwise. To determine statistical significance of differences between average metagene curves for two conditions, paired Wilcoxon signed rank tests were performed for each bin comparing normalized coverage values for each gene for this bin between the two conditions. P-values were adjusted for multiple testing with the Bonferroni method across all bins within each subfigure and are color-coded in the bottom track of subfigures: red = adj. p-value ≤ 10^− 15^, orange = adj. p-value ≤ 10^−10^, yellow = adj. p-value ≤ 10^−3^.

For hierarchical clustering analysis, PRO-seq profiles for each gene and condition were calculated for sense or antisense strand as for metagene analyses (without averaging across genes). PRO-seq profiles in promoter windows for mock and WT-F infection at 3 h p.i. were then concatenated and divided by the maximum value in the concatenated vector. Hierarchical clustering was performed in using the hclust function in R according to Euclidean distances and Ward’s clustering criterion. Peaks in metagene plots for each cluster were then determined in the following way: First, all local and global maxima and minima of metagene curves for each condition were identified for each cluster using the find_peaks function in the R ggpmisc package. The major peak was the global maximum. Subsequently, the next highest local maxima up- or downstream of the major peak were determined and retained as secondary peaks if (i) they were sufficiently removed from the borders of the 6 kb promoter window (i.e., within bins 30 to 80 of the 101 bins), (ii) the difference between the height of the secondary peak and the minimum value between the major and secondary peak was at least 10% of the major peak height and (iii) the height of the secondary peak was at least 20% of the major peak height.

### Over- and under-representation analysis

Over- and under-representation of Gene Ontology (GO) terms and transcription factor binding motifs from TRANSFAC was performed for each cluster using the g:Profiler webserver (59) and the R package gprofiler2 (60), which provides an R interface to the webserver. P-values were corrected for multiple testing using the Benjamini-Hochberg False Discovery Rate (FDR) (61) and significant terms or motifs were identified at an adjusted p-value cutoff of 0.001.

### Calculation of GC content and GC skew

Genome sequences in the ± 3kb around the TSS for each gene were extracted from the hg19 genome with twoBitToFa (http://genome.ucsc.edu/goldenPath/help/twoBit.html) and mean GC content and GC skew (G-C/G+C) was calculated in 100 bp sliding windows with steps of 1 bp as described by Watts *et al*. (34).

### Quantification of read-in transcription

Number of fragments (=read pairs) per gene or in the 5 kb upstream of a gene were determined from mapped paired-end 4sU-seq reads in a strand-specific manner using featureCounts (56) and gene annotations from Ensembl (version 87 for GRCh37). For genes, all fragments overlapping exonic regions on the corresponding strand by ≥ 25bp were counted for the corresponding gene. For the 5 kb upstream regions, all fragments overlapping the 5 kb upstream of the gene 3’end were counted. Gene expression and upstream transcriptional activity were quantified in terms of fragments per kilobase of exons per million mapped reads (FPKM). Only reads mapped to the human genome were counted for the total number of mapped reads for FPKM calculation. Percentage of read-in transcription was calculated as previously described (7, 39) for 7,271 genes that had no up- or downstream gene within 5□kb and were well expressed (average FPKM over replicates ≥ 1) in at least one time point of our 4sU-seq time-course. For this purpose, the percentage of transcription upstream of a gene was first calculated separately for each replicate as: Percentage of upstream transcription = 100 × (FPKM in 5 kb upstream of gene)/(gene FPKM), and averaged between replicates. Second, percentage of read-in at each 4sU-seq time-point of infection was calculated as percentage of upstream transcription in infected cells minus the percentage of downstream transcription in uninfected cells. Negative values were set to 0.

### Code availability

Workflows for PI calculation, metagene analyses, clustering and figure creation were implemented and run in Watchdog (47, 48) and are available at https://doi.org/10.5281/zenodo.7322848. Corresponding Watchdog modules are available in the Watchdog module repository (https://github.com/watchdog-wms/watchdog-wms-modules/).

## Supporting information

Data Set S1

Data Set S2

## Acknowledgements

This work was funded by the Deutsche Forschungsgemeinschaft (DFG, German Research Foundation) in the framework of the Research Unit FOR5200 DEEP-DV (443644894) project FR 2938/11-1 and by grants FR2938/9-1 to C.C.F. and LD1275/6-1 to L.D.

