## Supplemental Figures for "HSV-1 infection induces a downstream shift of promoter-proximal pausing for most host genes"

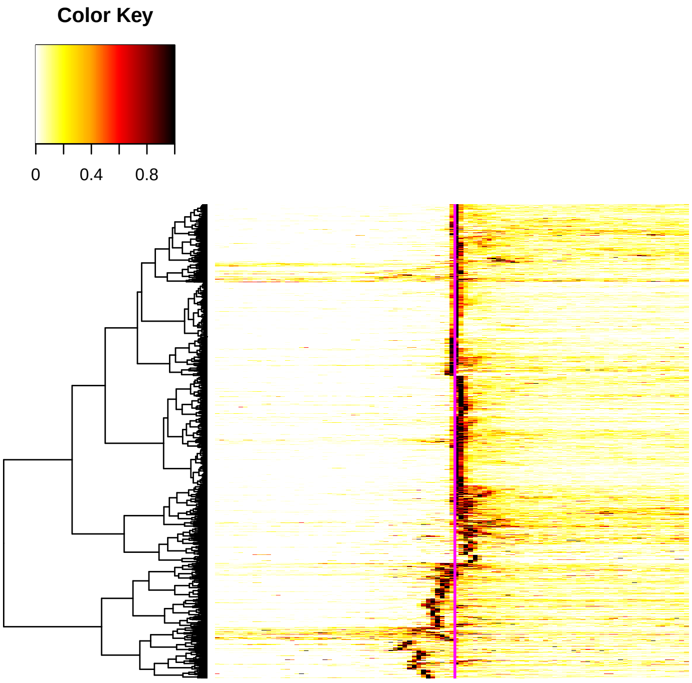


(a)

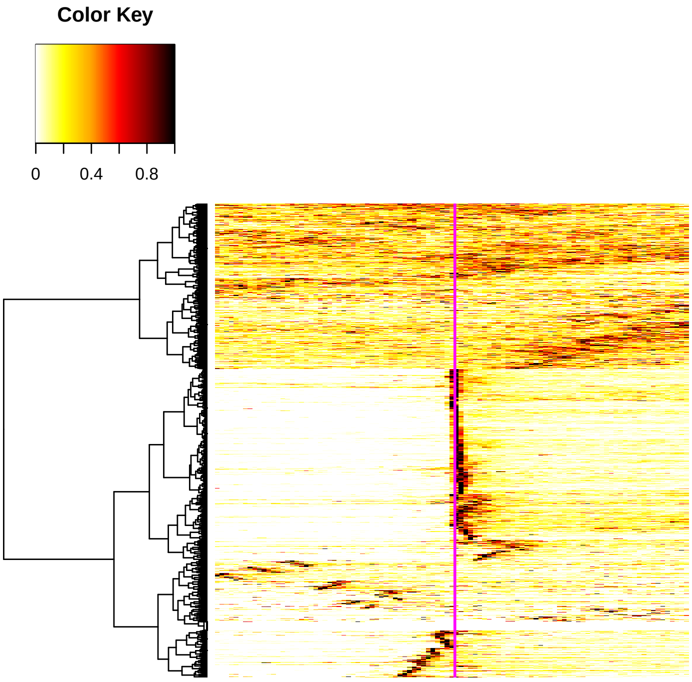


(b)

**Fig. S1** Heatmaps of PRO-seq profiles on the sense strand in mock infection in a window of ±3 kb around **(a)** the TSS positions identified from PROcap-seq and PRO-seq data of flavopiridol-treated HFF or **(b)** annotated gene 5' ends. For this purpose, PRO-seq profiles were divided by the maximum value in the ±3 kb promoter window, resulting in a value of 1 for the position of the highest peak in PRO-seq profiles. Hierarchical clustering of normalized PRO-seq profiles for all genes was performed using the *hclust* function in R according to Euclidean distances and Ward's clustering criterion. The central position in the promoter window (= the identified TSS) is marked by a vertical magenta line.

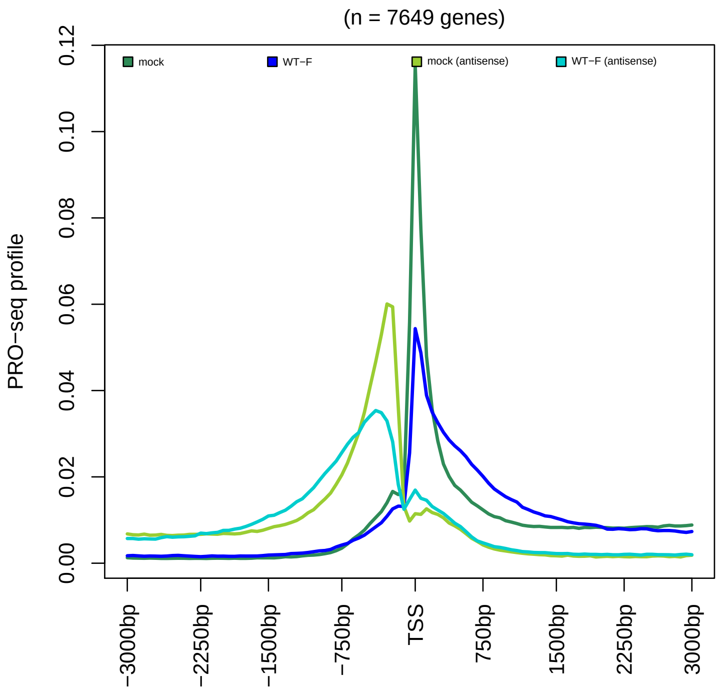


**Fig. S2** Metagene plot showing the distribution of PRO-seq profiles in sense (dark green and blue) and antisense (light green and blue) direction from -3 kb to +3 kb around the TSS for all analyzed genes for mock infection (dark and light green) and WT-F 3 h p.i. infection (dark and light blue). One gene without reads in any of the analyzed samples was excluded.

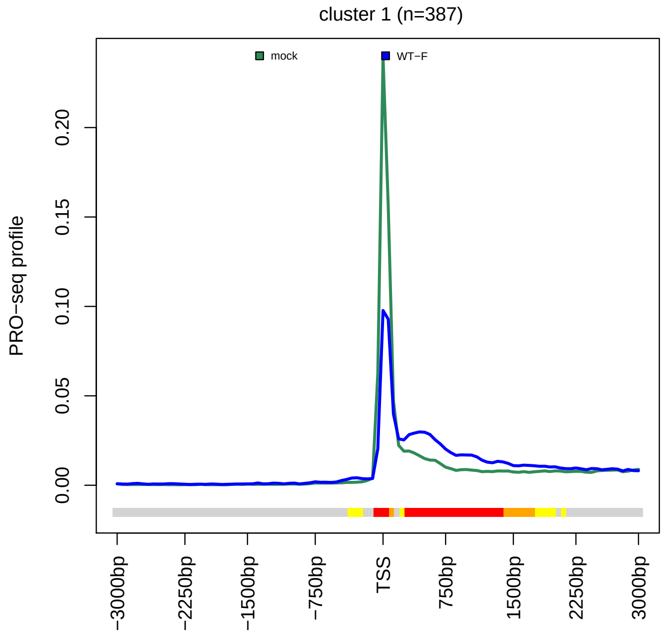


(a)

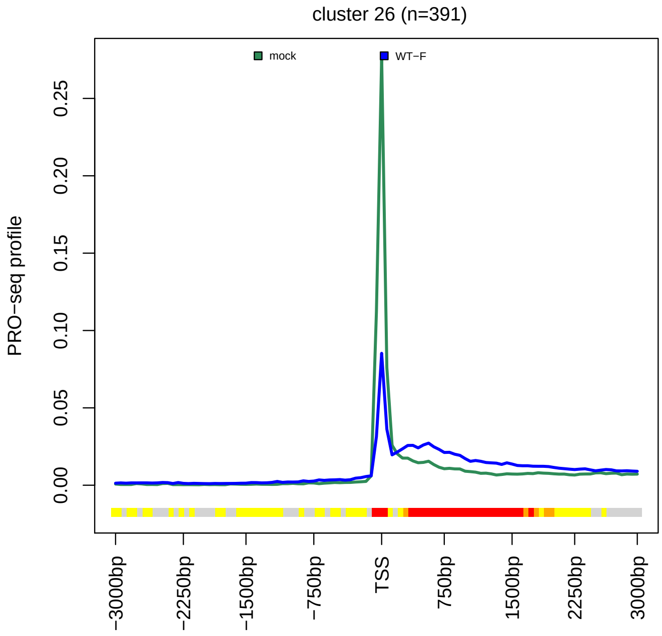


(b)

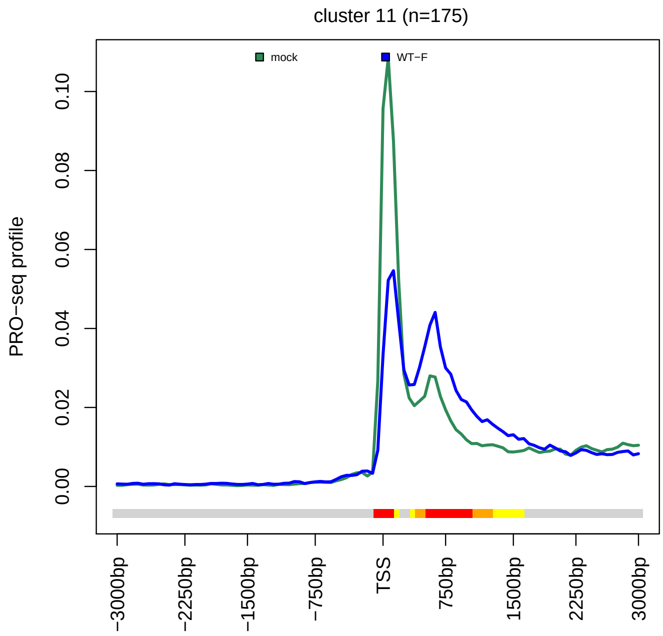


(c)

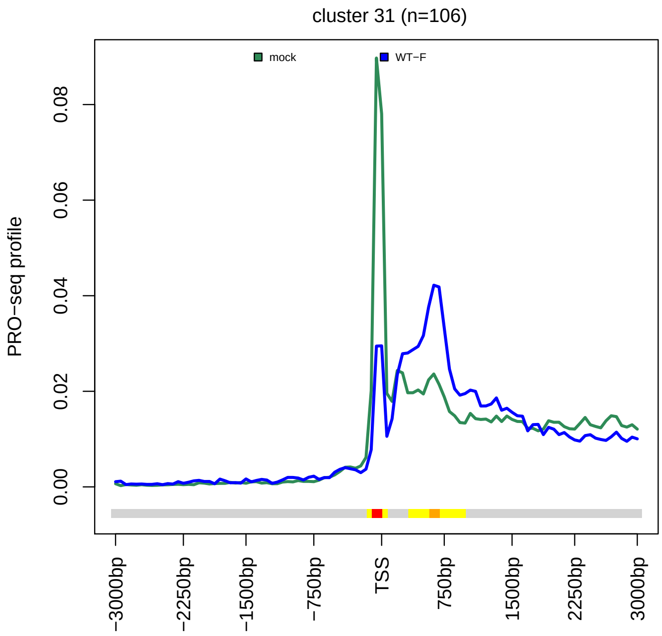


(d)

(Continued on next page)

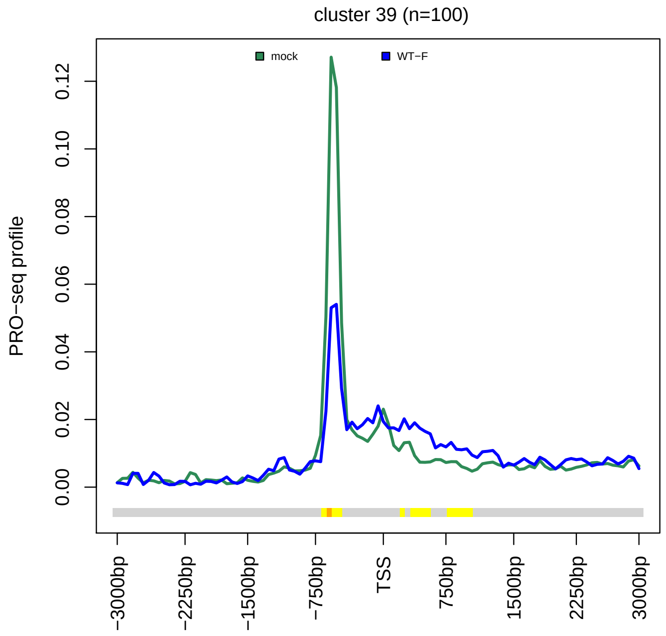


**(e)**

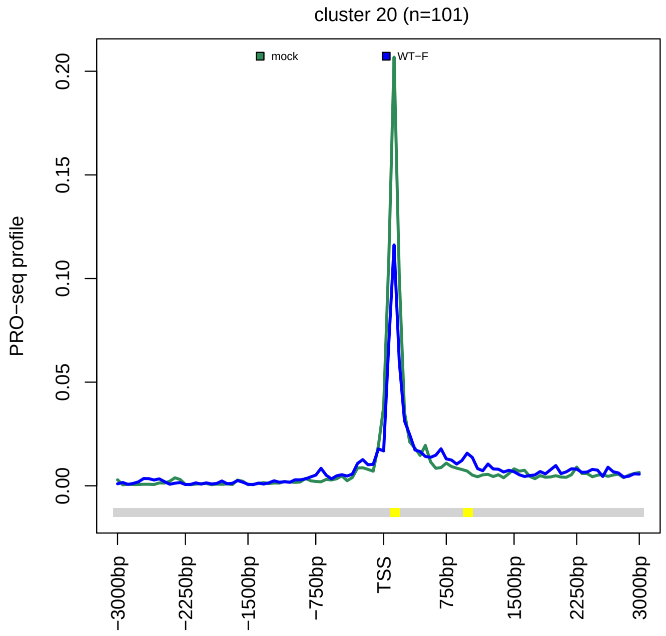


**(f)**

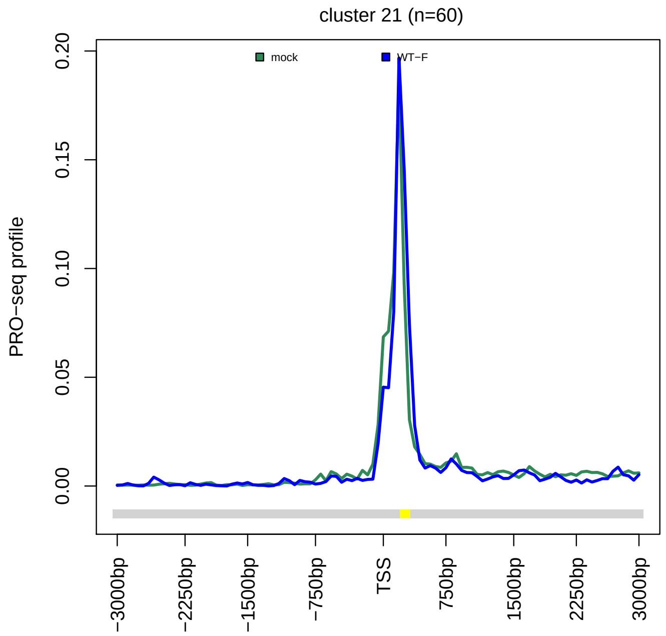


**(g)**

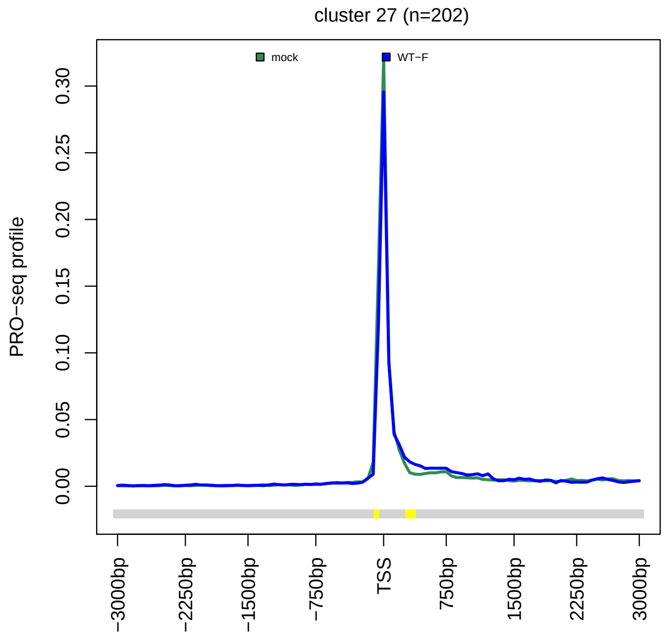


**(h)**

**Fig. S3** Metagene plots showing the PRO-seq profile in sense direction from -3 kb to +3 kb around the TSS for mock infection (dark green) and WT-F 3 h p.i. infection (dark blue) separately for example clusters. Cluster numbers and number of genes in each cluster are indicated on top of subfigures. The color track at the bottom of each subfigure indicates the significance of paired Wilcoxon tests comparing the normalized PRO-seq coverages of genes for each bin between mock and WT-F 3 h p.i. infection. P-values are adjusted for multiple testing with the Bonferroni method within each subfigure; color code: red = adj. p-value $\leq 10^{-15}$, orange = adj. p-value $\leq 10^{-10}$, yellow = adj. p-value $\leq 10^{-3}$.

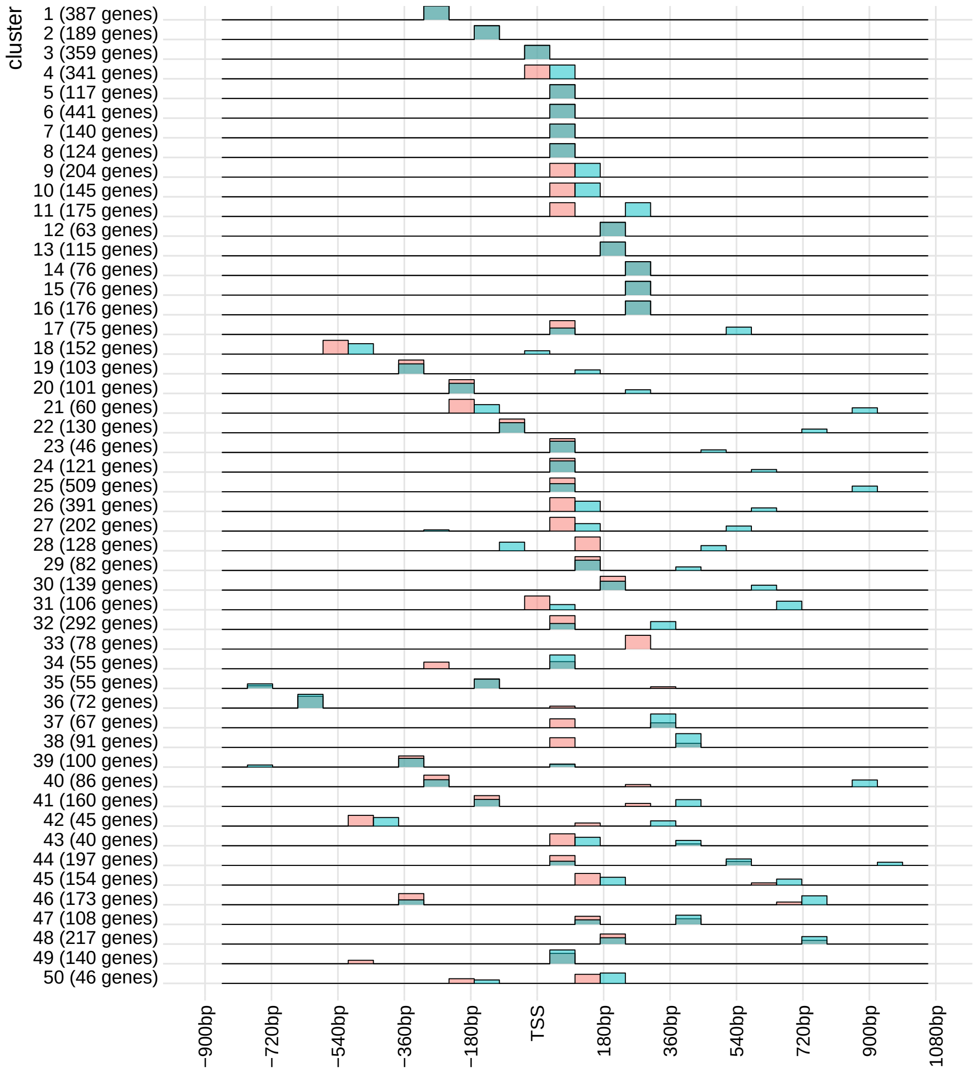


(a)

| Peak pattern | mock no. clusters | mock no. genes | WT-F 3 h p.i. no. clusters | WT-F 3 h p.i. no. genes |
|---|---|---|---|---|
| One TSS peak | 33 | 5728 | 21 | 3347 |
| Additional minor peak downstream of major TSS peak | 13 | 1461 | 17 | 2986 |
| Two approximately equally high peaks | 2 | 232 | 4 | 296 |
| Additional downstream peak higher than TSS peak | 0 | 0 | 7 | 834 |
| none of the above | 2 | 228 | 1 | 46 |

(b)

**Fig. S4** **(a)** Positions, number, and relative heights of peaks identified in PRO-seq profiles in sense direction for the 50 clusters. Mock infection is shown in light red and WT-F 3 h p.i. infection in turquoise. Darker turquoise indicates that a peak is present at the same position in mock and WT-F 3 h p.i. infection. The relative peak height is calculated as the peak height divided by the sum of all peak heights for the same condition. Thus, a single peak has a value of 1, two equally high peaks both have a value of 0.5, and so on. **(b)** Statistics on the number of clusters and number of genes with different types of peak patterns defined by the number and relative height of peaks for mock and WT-F 3 h p.i. infection shown in **(a)**.

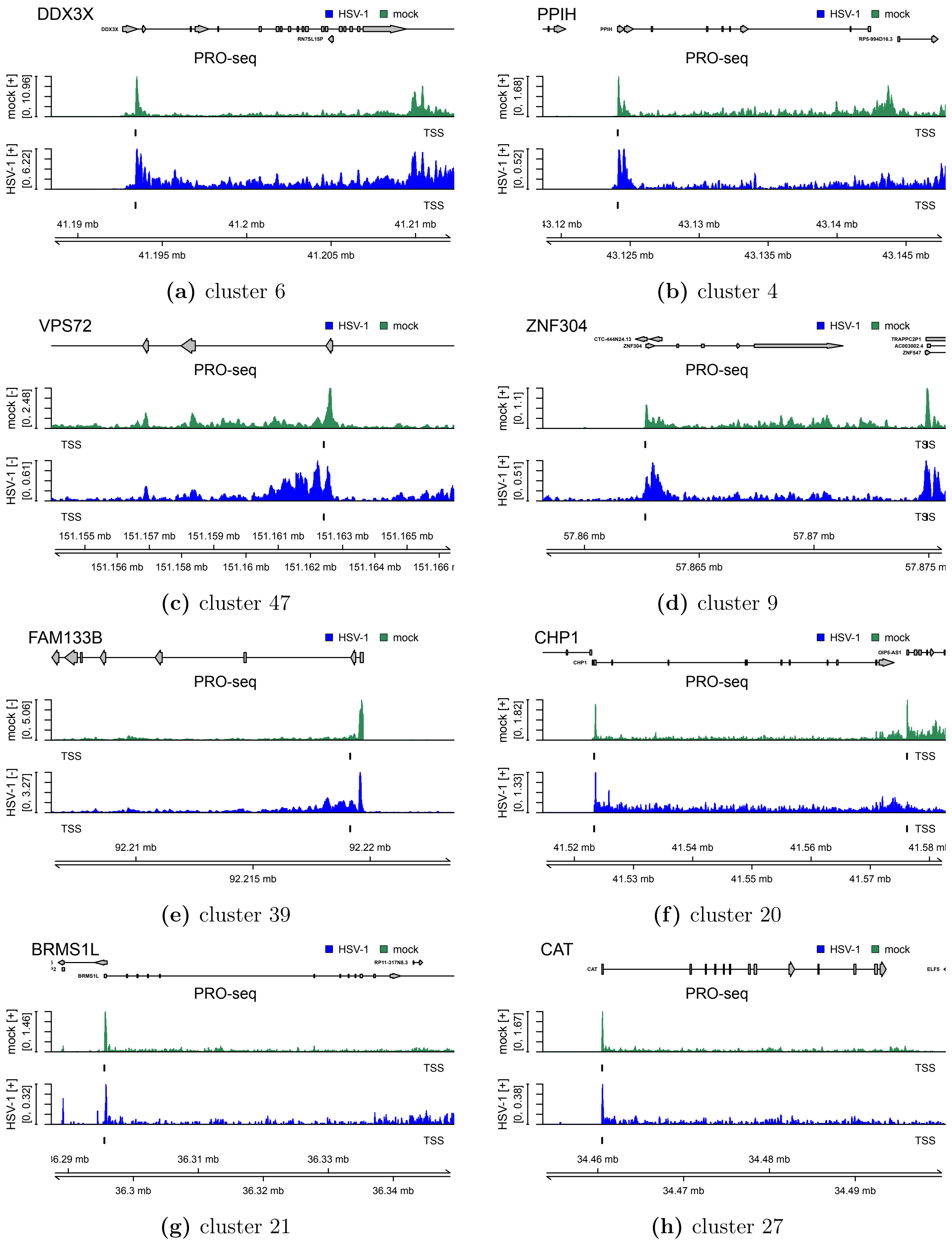


**(a)** cluster 6

**(b)** cluster 4

**(c)** cluster 47

**(d)** cluster 9

**(e)** cluster 39

**(f)** cluster 20

**(g)** cluster 21

**(h)** cluster 27

**Fig. S5** Read coverage around the TSS in PRO-Seq data (sense strand only) for mock (green) and WT-F infection (blue) at 3 h p.i. for example genes (gene name of the selected gene on the top left) in different clusters (cluster number shown below subfigures). Read coverage was normalized to total number of mapped reads and averaged between replicates. The identified TSS used in the analysis is indicated by a short vertical line below each read coverage track. Gene annotation is indicated at the top. Boxes represent exons, lines represent introns and direction is indicated by arrowheads. Genomic coordinates are shown on the bottom. Please note that figures are not centered around the TSS, but a larger region downstream of the TSS was included than upstream of the TSS.

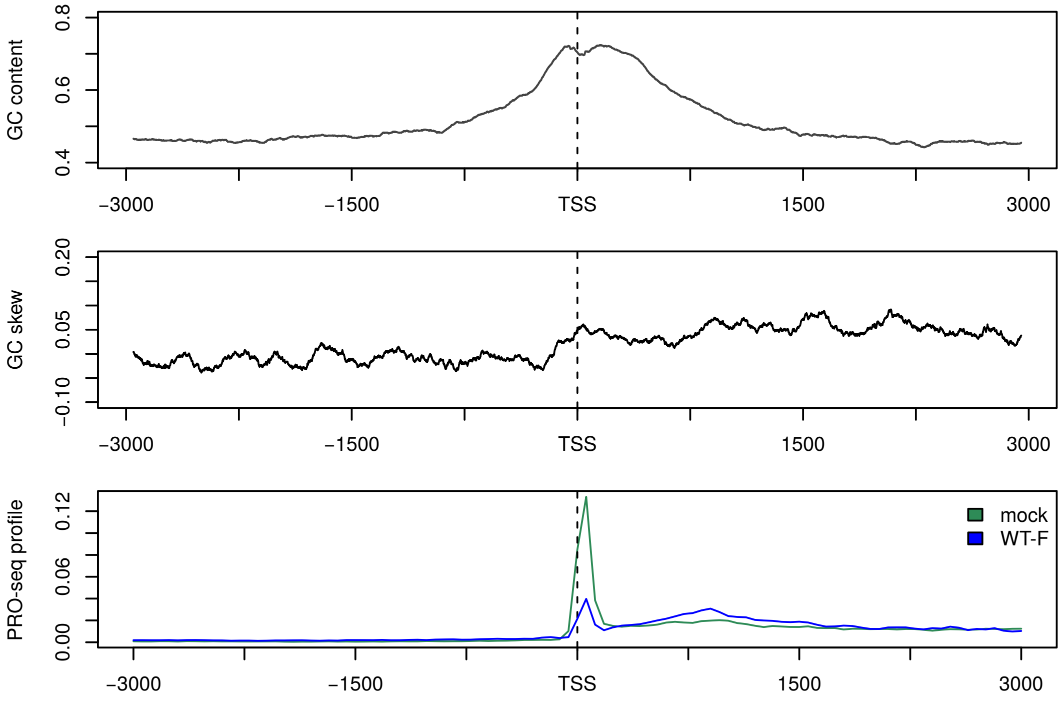


**(a)** cluster 32

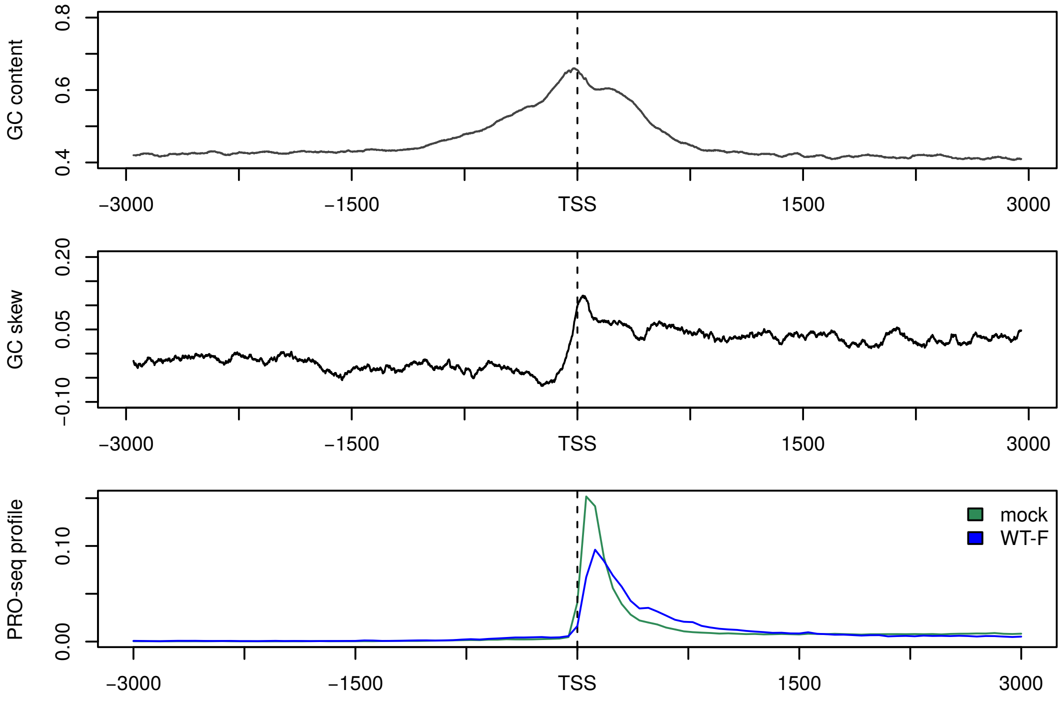


**(b)** cluster 6

**(Continued on next page)**

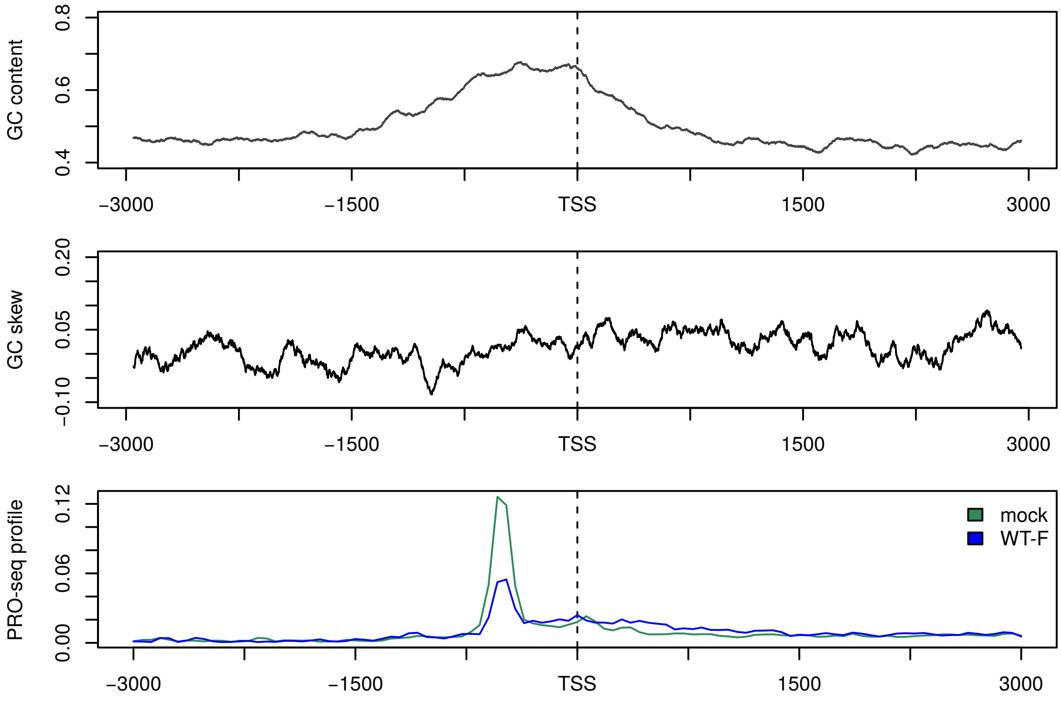


**(c)** cluster 39

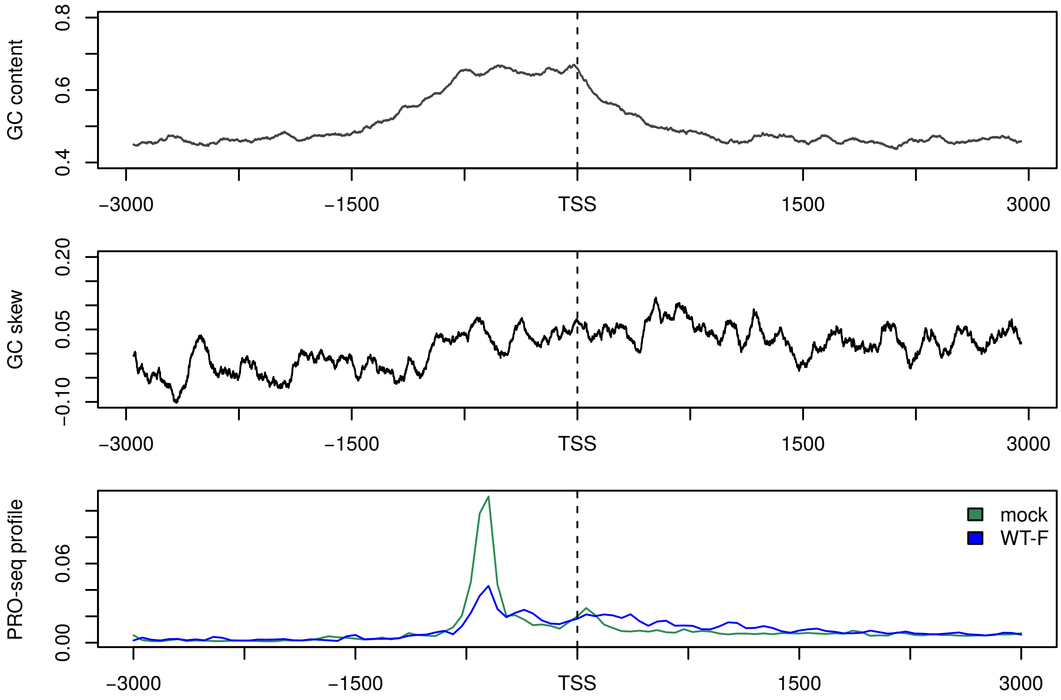


**(d)** cluster 40

**Fig. S6** GC content and GC skew in promoter regions for example clusters (cluster numbers shown beneath subfigures). For each gene GC content and GC skew was determined in 100 bp sliding windows from -3 kb of the TSS to +3 kb of the TSS. Values for each sliding window were then averaged across genes in this cluster. The bottom panel of each subfigure shows the PRO-seq profiles in mock and WT-F 3 h p.i. infection for comparison.

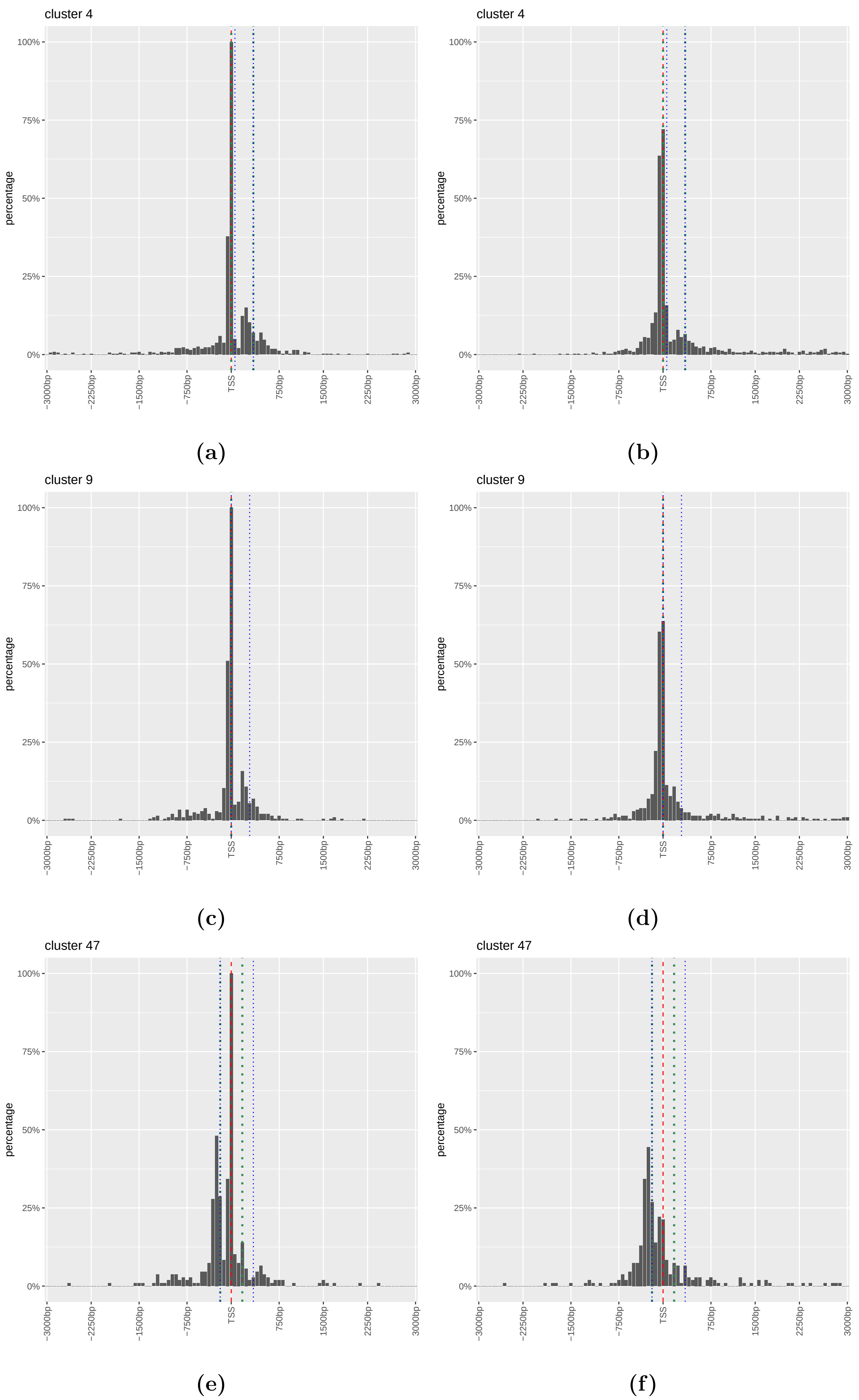


**Fig. S7 (a,c,e)** Percentage of genes exhibiting a peak in the PROcap-seq and PRO-seq data of flavopiridol-treated HFF at particular positions for example clusters (indicated on the top left of subfigures). This includes all identified peaks for a gene not just the major peak used for identifying the TSS. For this purpose, the region ± 3 kb around the identified peak was divided into bins of 60 bp and for each bin the percentage of genes with a peak falling into this bin were calculated. The red dashed vertical line marks the identified TSS. Green dotted vertical lines indicate peak positions in mock infection and blue dotted vertical lines peak positions in WT-F infection at 3 h p.i. **(b,d,f)** Percentage of genes exhibiting an annotated TSS in each 60 bp bin around the identified TSS for example clusters (indicated on the top left of subfigures). TSS and peak positions in WT-F infection at 3 h p.i. are indicated as in **(a,c,e)**.

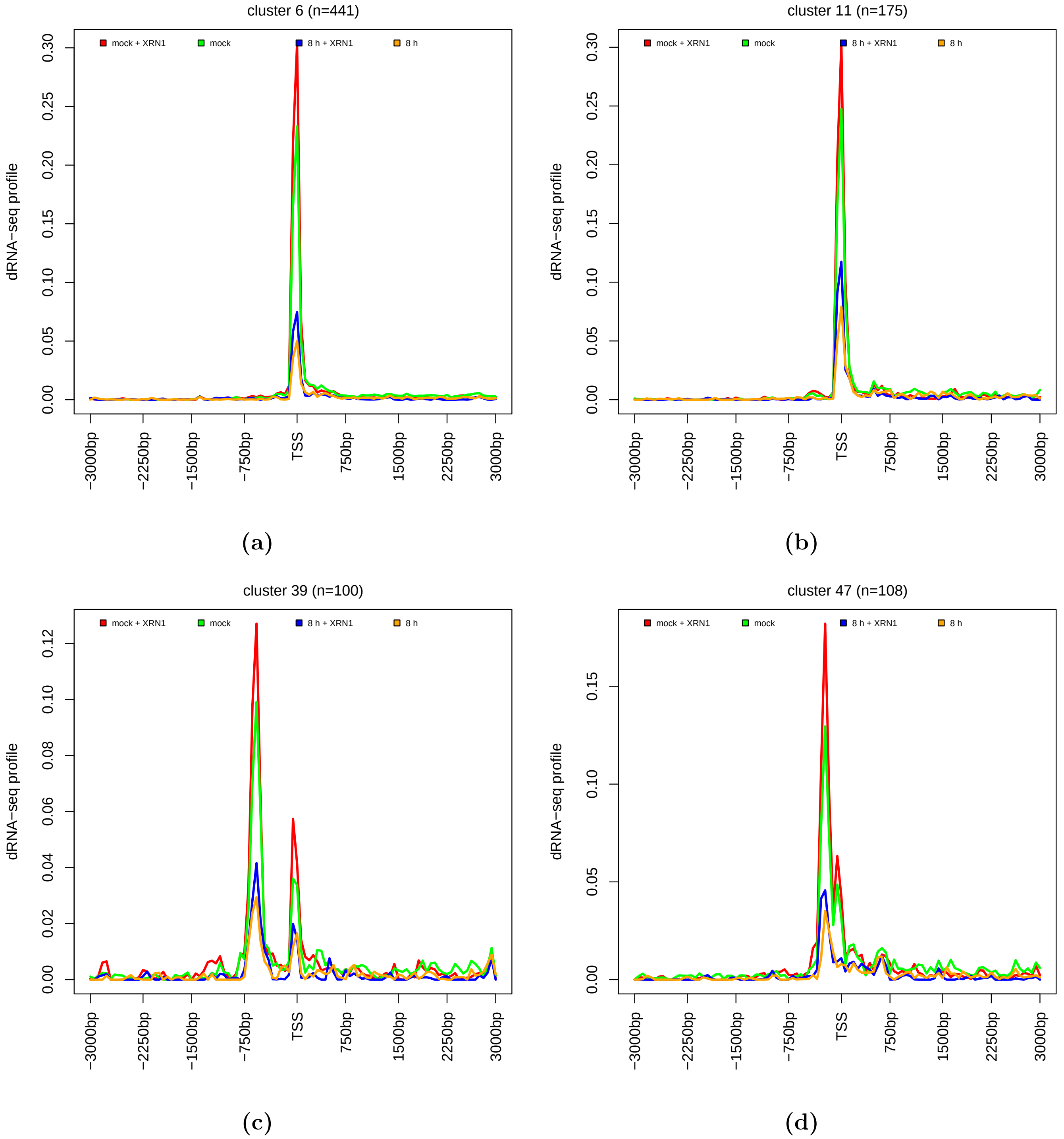


**Fig. S8** Metagene plots of dRNA-seq profiles on the sense strand in mock and WT-17 8 h p.i. infection with and without XRN1 treatment for example Clusters 6, 11, 39, and 47, which show broadening of peaks or additional peaks originating or increasing in height in PRO-seq data during WT-F infection. For metagene plots of PRO-seq profiles for these clusters see Fig. 2 and Fig. S3.

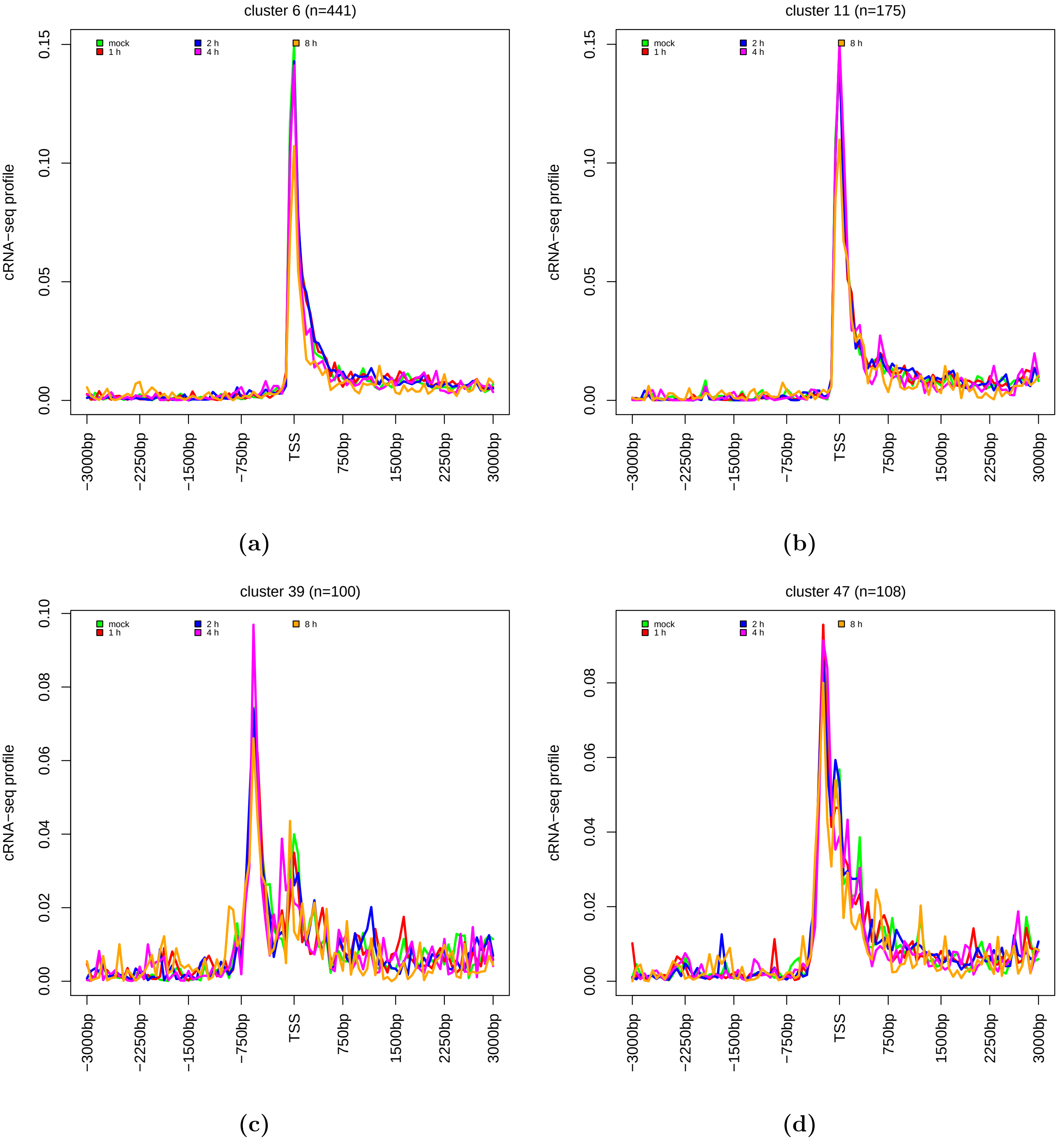


**Fig. S9** Metagene plots of cRNA-seq profiles on the sense strand in mock and WT-17 infection at 1, 2, 4, and 8 h p.i. for example Clusters 6, 11, 39, and 47, which show broadening of peaks or additional peaks originating or increasing in height in PRO-seq data during WT-F infection. For metagene plots of PRO-seq profiles for these clusters see Fig. 2 and Fig. S3.

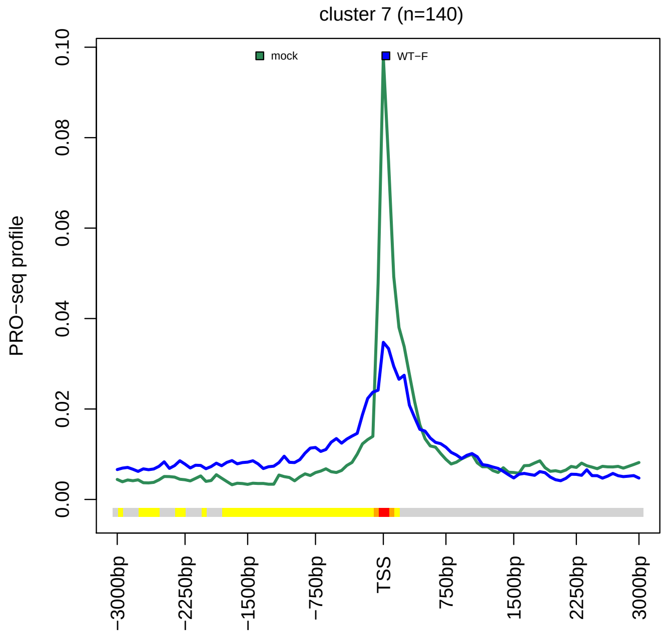


(a)

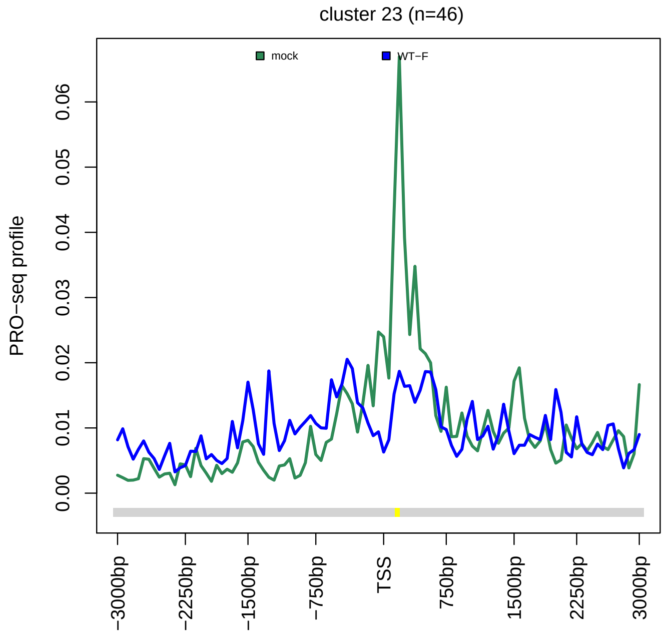


(b)

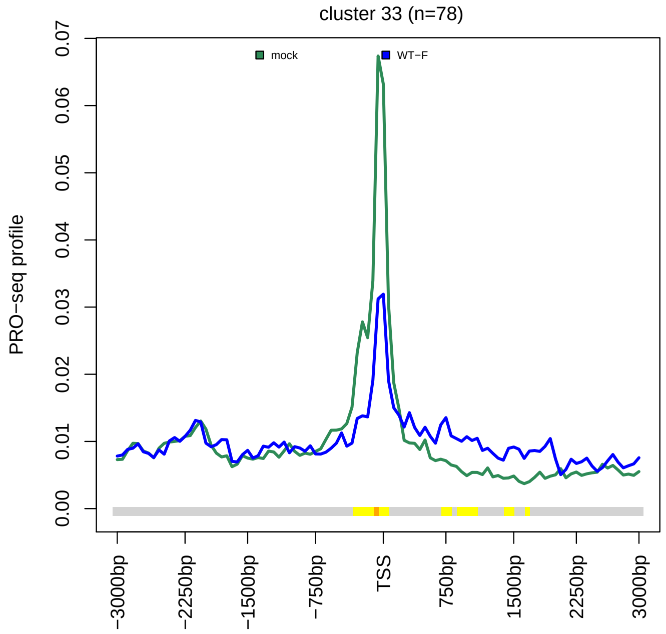


(c)

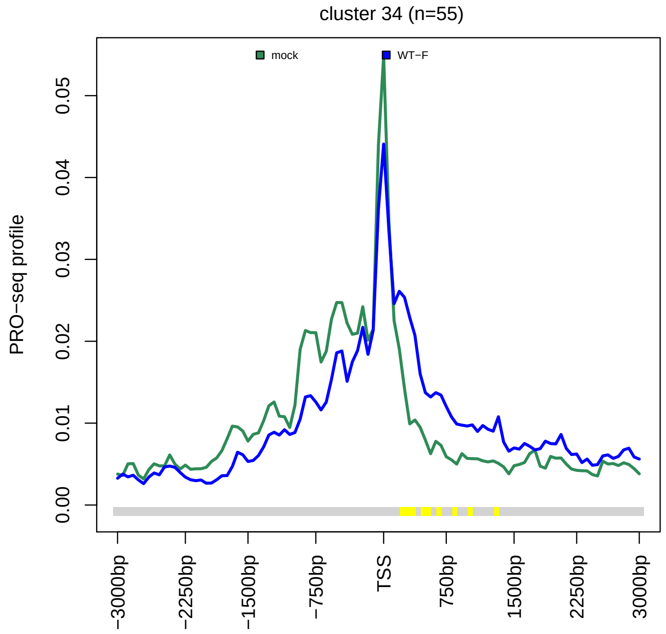


(d)

**(Continued on next page)**

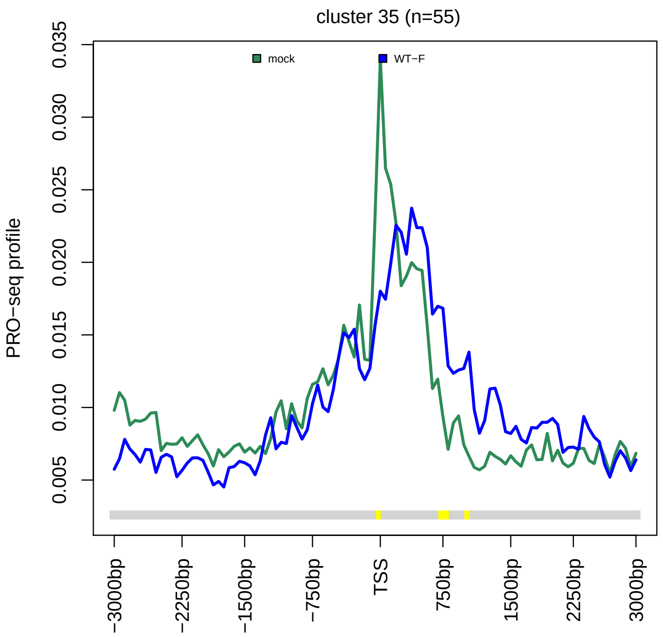


**(e)**

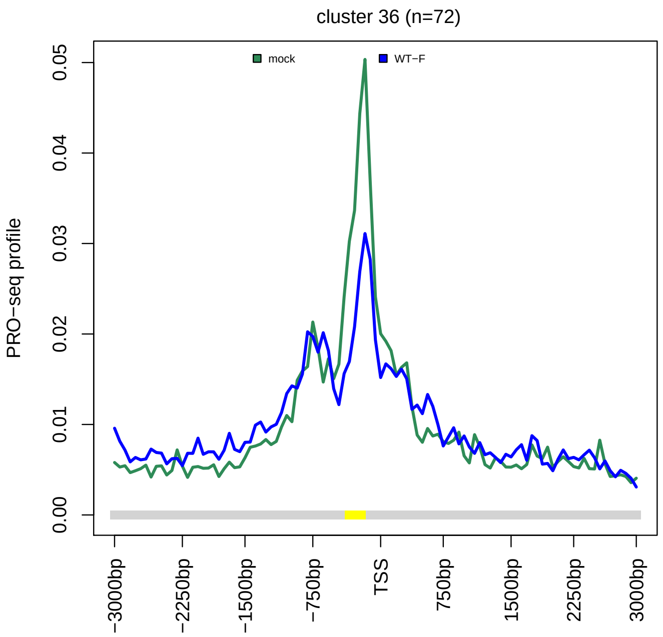


**(f)**

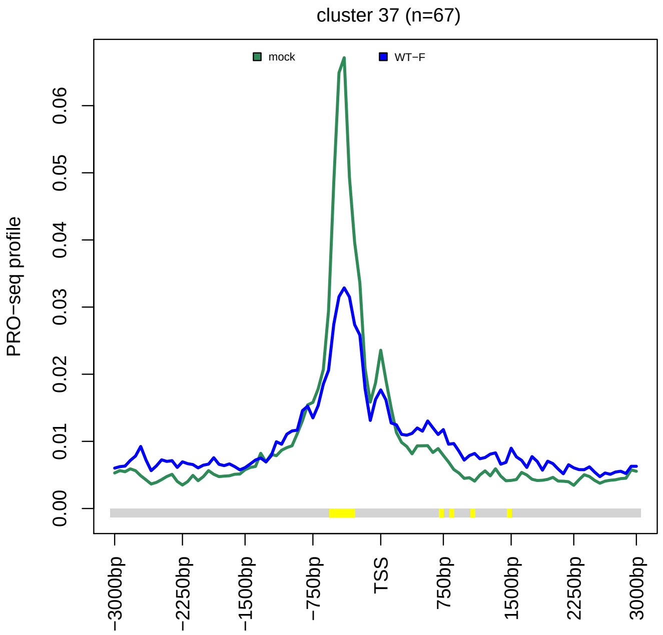


**(g)**

**Fig. S10** Metagene plots showing the PRO-seq profile in sense direction from -3 kb to +3 kb around the TSS for mock infection (dark green) and WT-F 3 h p.i. infection (dark blue) separately for Clusters 7, 23, and 33 to 37, which exhibit a small extent of read-in transcription in 3-4 h p.i. 4sU-seq (see Fig. 4). Cluster numbers and number of genes in each cluster are indicated on top of subfigures. The color track at the bottom of each subfigure indicates the significance of paired Wilcoxon tests comparing the normalized PRO-seq coverages of genes for each bin between mock and WT-F 3 h p.i. infection. P-values are adjusted for multiple testing with the Bonferroni method within each subfigure; color code: red = adj. p-value $\leq 10^{-15}$, orange = adj. p-value $\leq 10^{-10}$, yellow = adj. p-value $\leq 10^{-3}$.

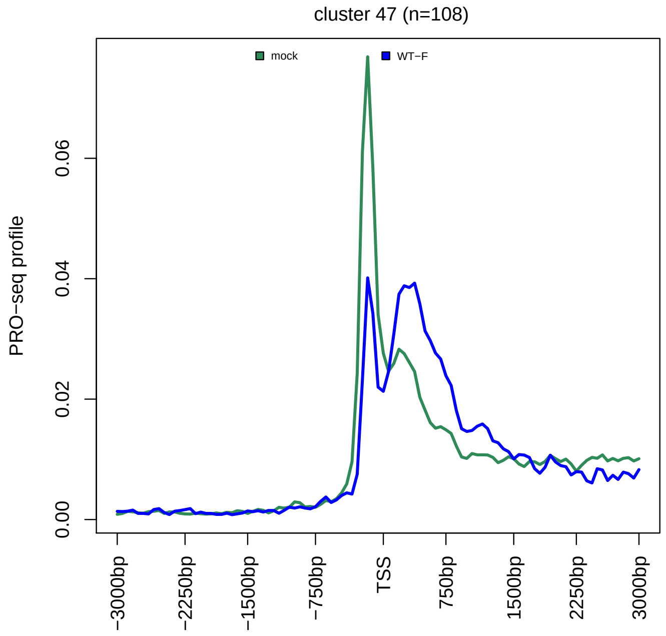


(a)

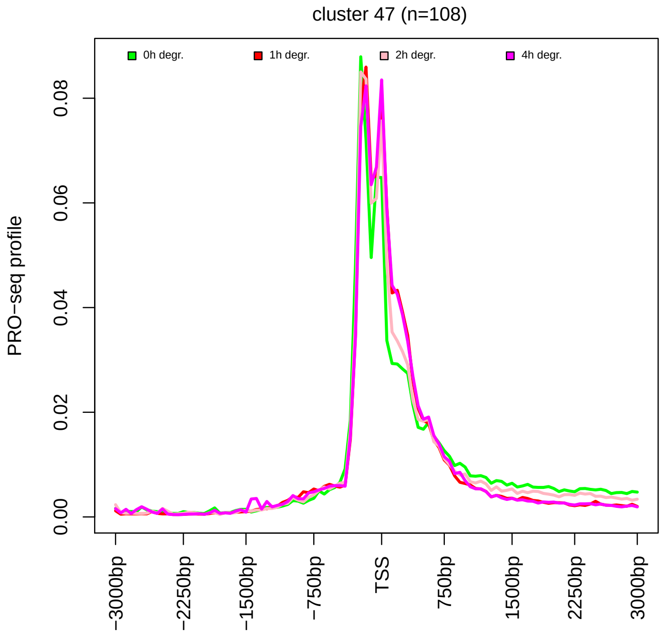


(b)

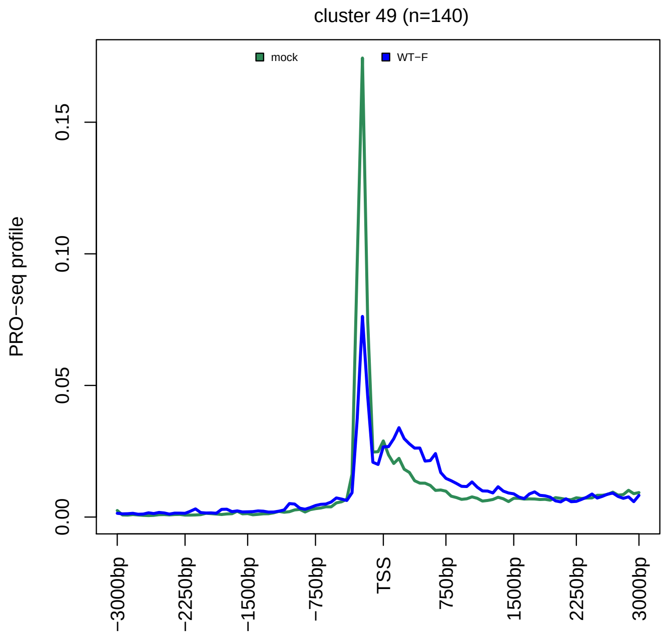


(c)

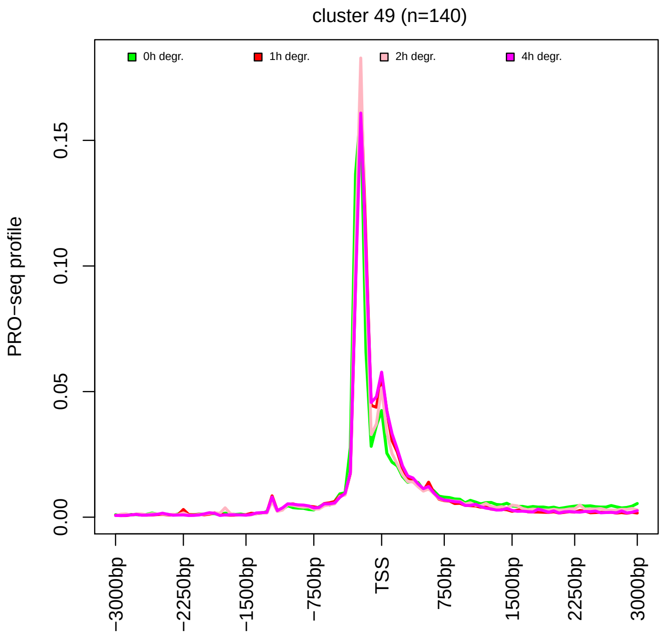


(d)

(Continued on next page)

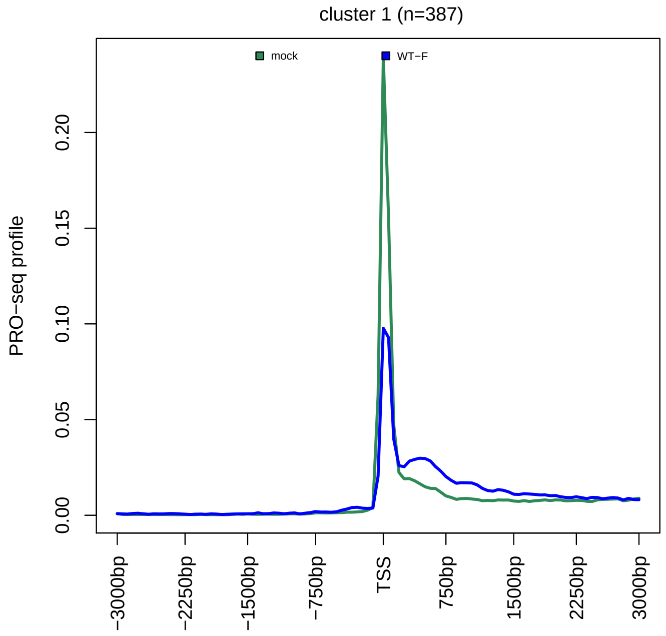


(e)

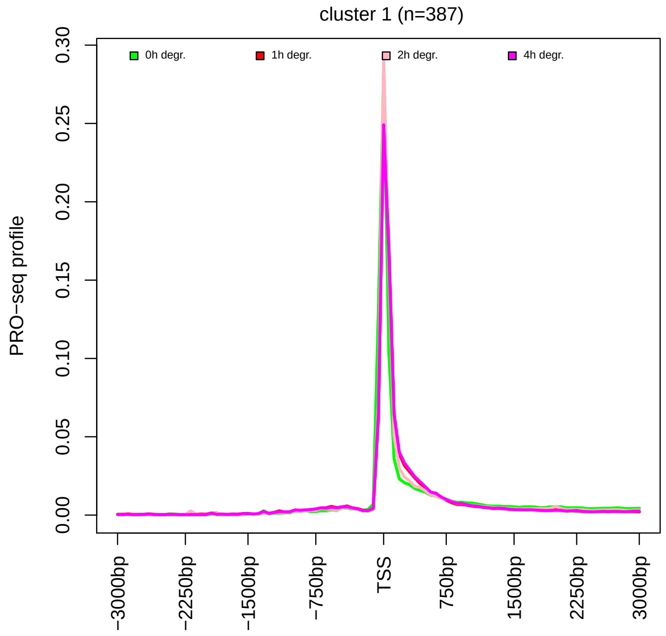


(f)

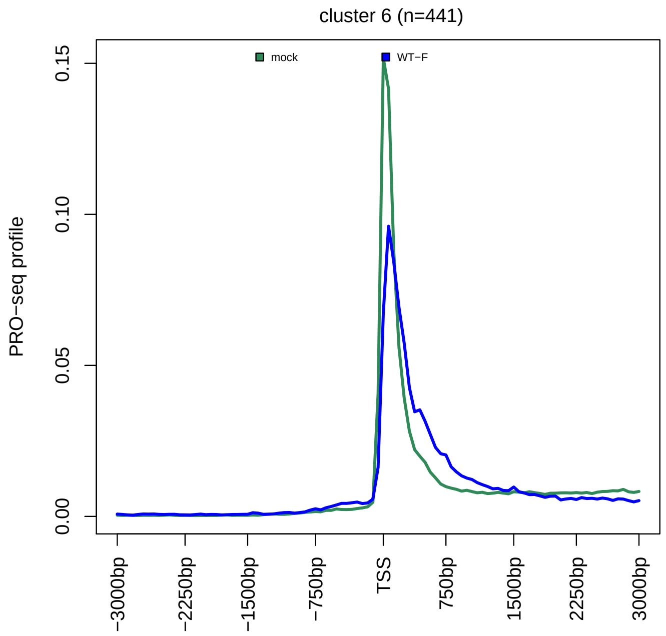


(g)

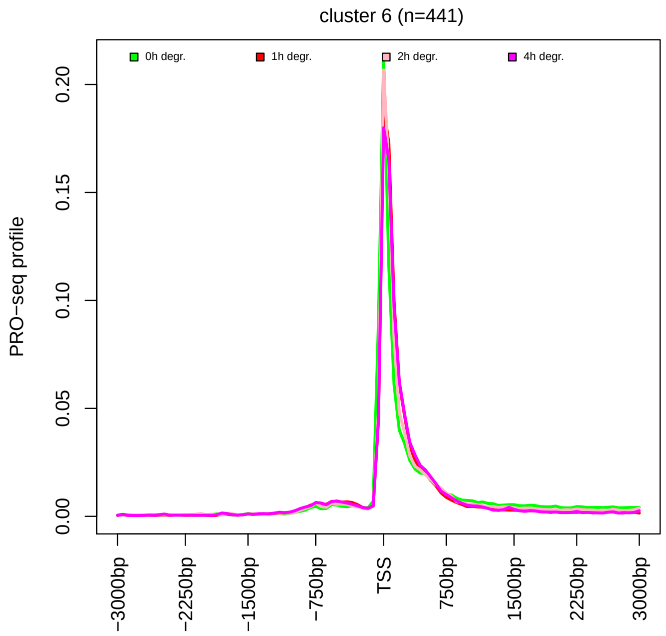


(h)

(Continued on next page)

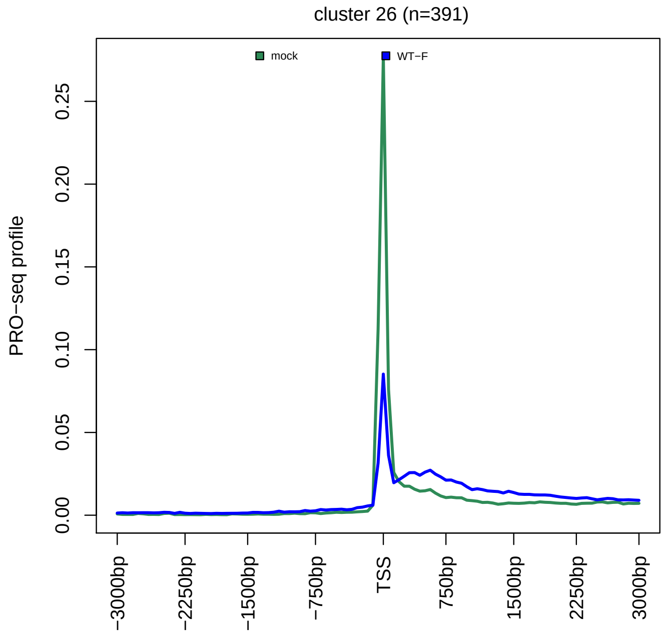


(i)

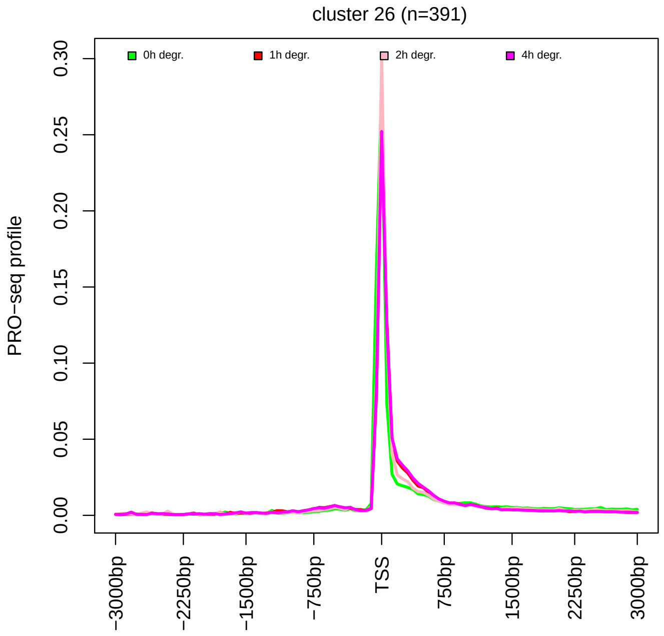


(j)

**Fig. S11** Metagene plots around the TSS of PRO-Seq profiles for mock and WT-F 3 h p.i. infection from the study of Birkenheuer *et al.* (left column) and 0, 1, 2, and 4 h auxin-inducible degradation of NELF from the study by Aoi *et al.* (right column) for example clusters showing **(a-d)** an increased downstream peak or **(e-j)** only a reduced and slightly broadened TSS peak upon NELF degradation.
